# From iPSCs to NPCs to cortical neurons: bioenergetic, neuronal, and calcium signaling phenotypes in bipolar disorder with and without familial mitochondrial disease

**DOI:** 10.64898/2025.12.19.695603

**Authors:** Dana El Soufi El Sabbagh, Jaehyoung Choi, Lakshmini Balachandar, Eileen Haydarian, Jude Demyati, Zaara Mazhar, Angela Duong, Chiara Cristina Bortolasci, Courtney Irwin, Karun K Singh, Michael Berk, Ken Walder, L Trevor Young, Peter L. Carlen, Ana Cristina Andreazza

**Affiliations:** Department of Pharmacology and Toxicology, University of Toronto, Toronto, ON, Canada; Mitochondrial Innovation Initiative, MITO2i, Toronto, ON, Canada; Krembil Brain Institute, Toronto Western Hospital, University Health Network, Toronto, ON, Canada; Department of Physiology, University of Toronto, Toronto, ON, Canada; Deakin University, IMPACT - the Institute for Mental and Physical Health and Clinical Translation, School of Medicine, Barwon Health, Geelong, Australia; Donald K. Johnson Eye Institute, Krembil Research Institute, University Health Network, Toronto, ON, Canada; Department of Laboratory Medicine and Pathobiology, Faculty of Medicine, University of Toronto, Toronto, ON, Canada; Center for Addiction and Mental Health, Toronto, ON, Canada; Department of Psychiatry, University of Toronto, Toronto, ON, Canada

**Author notes:** Correspondent Author: Ana C. Andreazza University of Toronto, 777 Bay Street, Room 1031. Toronto, ON, M5G 2R3.

**Keywords:** Bipolar Disorder, mitochondrial dysfunction, iPSC, hyperexcitability, mitochondrial disease, cortical neurons

## Abstract

**Background:** Psychiatric disorders frequently accompany primary mitochondrial diseases (PMDs), implicating mitochondrial dysfunction as a shared biological substrate for psychiatric vulnerability. To determine how familial mitochondrial risk influences neuronal development in bipolar disorder (BD), we generated induced pluripotent stem cells (iPSCs), neural progenitor cells (NPCs), and cortical neurons (CNs) from three healthy controls (CT), three patients with BD, and three patients with BD and a family history of mitochondrial disease (BD-FMD).

**Methods:** Across differentiation, we assessed mitochondrial function (ATP production, mitochondrial membrane potential, ROS generation, cytosolic cell-free mtDNA), metabolomic signatures, calcium imaging, and neuronal electrophysiological activity through multi-electrode arrays (MEA).

**Results:** BD neurons uniquely exhibited pronounced hyperexcitability in comparison to CT and BD-FMD groups. The BD-FMD group displayed prolonged mitochondrial calcium transients, altered membrane potential, and aberrant ROS, in comparison to CT and BD, consistent with a sustained energetic deficit. Metabolomic profiling revealed distinct pathway enrichments in BD and BD-FMD, indicating divergent bioenergetic adaptations to mitochondrial burden.

**Conclusions:** Our findings reveal that familial mitochondrial liability actively reshapes neuronal differentiation and function, producing distinct trajectories of mitochondrial performance, calcium signaling, and network excitability. These results suggest that mitochondrial dysfunction in BD is not a secondary byproduct of illness, but a mechanistic contributor to altered neuronal energetics and communication. Together, these findings delineate how inherited mitochondrial vulnerability reshapes neuronal excitability and metabolism, revealing bioenergetic phenotypes that may inform precision stratification and therapeutic targeting in BD.

## Introduction

Bipolar disorder (BD) is increasingly recognized as a disorder of disrupted cellular energetics, where mitochondrial dysfunction may underlie its episodic neurobiology. Despite affecting ∼2% of the global population, BD remains diagnosed purely by clinical criteria, and current treatments only restore mood stability for a subset of patients^1–4^. Understanding how mitochondrial risk alters neuronal development could reveal new paths for mechanistic stratification and treatment development.

Modeling neuronal processes and the contribution of mitochondrial is not simple. However, advancements in induced pluripotent stem cell (iPSC) models are starting to unveil links. For instance, BD iPSC-derived neural progenitor cells (NPCs) show varying proliferation rates and calcium dysregulation, while iPSC-derived neurospeheres exhibited smaller neurosphere sizes in BD, likely reflecting premature differentiation and neurodevelopment^5,6^. We have previously demonstrated that BD cerebral organoids (COs) have decreased ATP levels and heightened susceptibility to NLRP3 inflammasome activation, reinforcing a convergence on bioenergetic-immune dysregulation^7^. Additionally, transcriptomic profiling of BD COs highlighted immune signalling alterations with neurotransmission deficits, strengthening the link between energy consumption and neural circuit formation^8^. Supporting evidence shows that mitochondria are key regulators in controlling NPC fate decisions during differentiation via OXPHOS/glycolysis, reactive oxygen species (ROS) signalling, and mitophagy pathways^9–11^. This framework depicts that mitochondrial vulnerabilities may perturb cell-level neuronal developmental trajectories and downstream neuronal phenotypes in a developing brain and might provide a window to understand early pathophysiology in BD that couples compromised bioenergetics with altered neural development.

Psychiatric disorders frequently accompany familial mitochondrial diseases (MD), suggesting that cellular energy systems and mood regulation share mechanistic substrates^12–17^. In a recent population-based cohort study in Canada, our group has demonstrated that co-prevalent mental health conditions are common in patients with primary mitochondrial diseases (PMD)^14^. Approximately 18% of patients with PMD had a comorbid diagnosis of a mental health condition along with highest healthcare utilization, underscoring mitochondria’s central role in brain function^14,15^.

To dissect how familial mitochondrial risk shapes neuronal development, we compared a unique sample of patients with bipolar disorder (BD) with or without familial mitochondrial disease (BD-FMD), and non-psychiatric controls (CT). We combined mitochondrial function assessments, metabolomic pathway signatures, and neuronal firing phenotypes analysis across iPSCs, NPCs, and cortical neurons (CNs) to understand risk-dependent mitochondrial trajectories that could inform precision stratification in BD (Figure 1).

**Figure 1.**
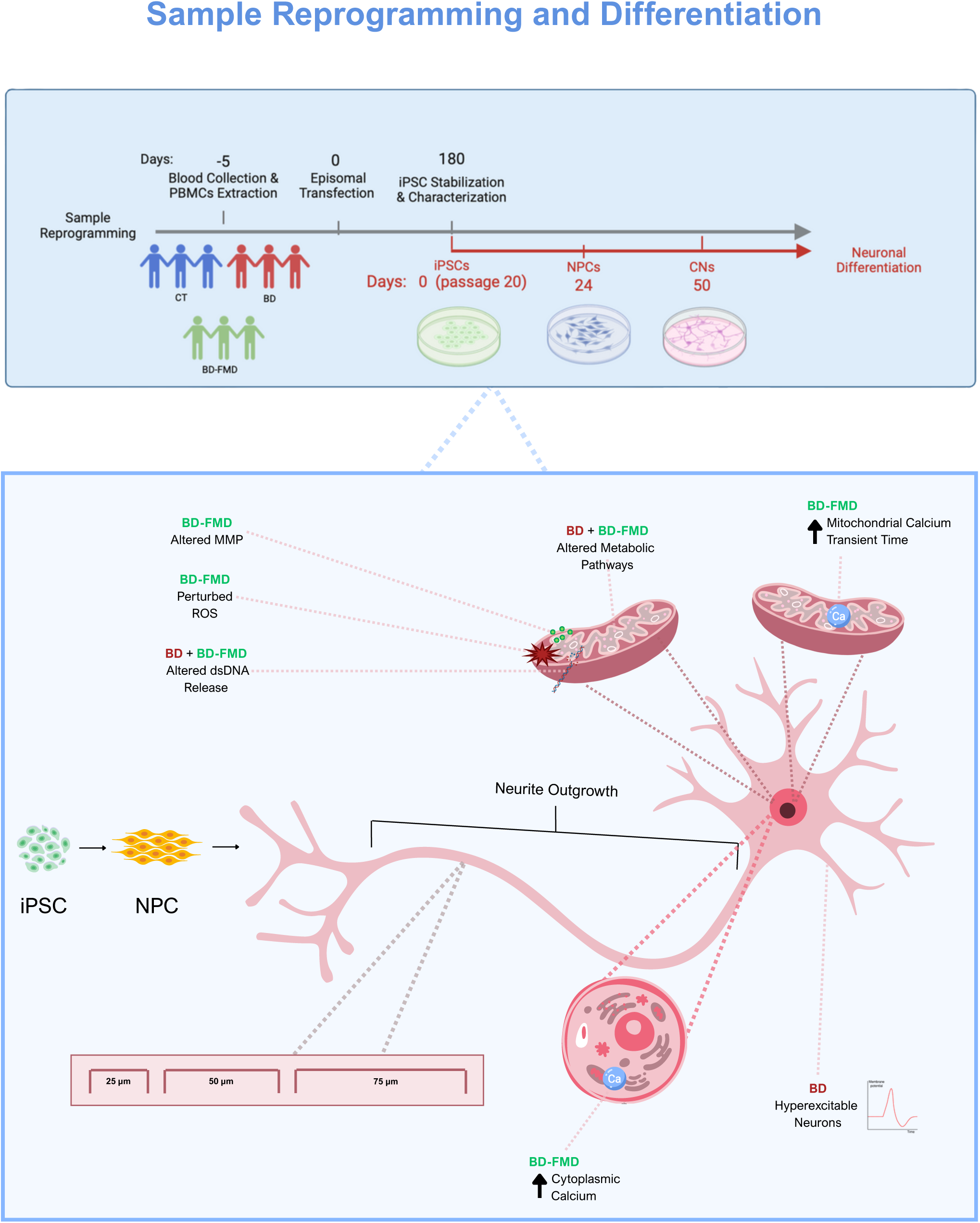
Overview of study design. Examining cellular health across iPSCs, NPCs, and CNs. Using the 3 disease groups (CT (blue), BD (red), BD-FMD (green). IPSCs, NPCs, and CNs were generated and underwent mitochondrial and cellular assays to investigate disease specific differences across neurodifferentiation timepoints.

## Methods and Materials

### Study Design

Nine participants were included in this study: three non-psychiatric controls (CT), three patients with bipolar disorder (BD), three patients with BD and a family history of clinically diagnosed mitochondrial disease (cytochrome c oxidase deficiency) (BD-FMD) (Figure 1).

All experimental protocols for this study were approved by the appointed research ethics board of University of Toronto Research ethics board (REB) (Protocol Number: 29949) and REB (Protocol Number: 36359) and at Deakin University with REB (17/205) in accordance with the Helsinki Declaration of 1975. Dr. Andreazza’s laboratory has also received approval from the Stem Cell Oversight Committee (Canadian Institutes of Health Research, #399222).

Generation, culture, karyotyping, and whole genome sequencing (WGS) of iPSC lines are described in Supplemental Methods.

### Differentiation of iPSC to Neural Progenitors and Cortical Neurons

IPSCs were differentiated into NPC using a dual-SMAD inhibition protocol and followed using the Forebrain Neuron Differentiation Kit (StemCell Technologies 08600) for CN generation. Full differentiation, characterization, intracellular and extracellular experiments involving ATP production, mitochondrial functional assessments, and metabolomic and lipidomic analyses are described in Tables 1-2 and the Supplementary Methods section.

**Table 1.** Antibodies used for immunofluorescence.

| Antibody | Type | Source | Identifier | Dilution |
| --- | --- | --- | --- | --- |
| MAP2 | Primary | Thermo Fisher | I3-500 | 1:200 |
| GFAP | Primary | Abcam | ab4674 | 1:500 |
| VGLUT1 | Primary | Synaptic Systems | 135303 | 1:200 |
| GABA | Primary | Sigma | A2052 | 1:200 |
| OCT4 | Primary | Invitrogen | PA5-27438 | 1:200 |
| NESTIN | Primary | Invitrogen | MA1-110 | 1:200 |
| Goat anti-mouse<br>CY3 | Secondary | Abcam | #Ab97935 | 1:750 |
| Donkey anti-rabbit Alexa<br>Fluor Plus 647 | Secondary | Invitrogen | #A32795 | 1:750 |
| Donkey anti-chicken 488 | Secondary | Jackson<br>ImmunoResearch<br>Laboratories Inc | #703-546-155 | 1:750 |

**Table 2.**
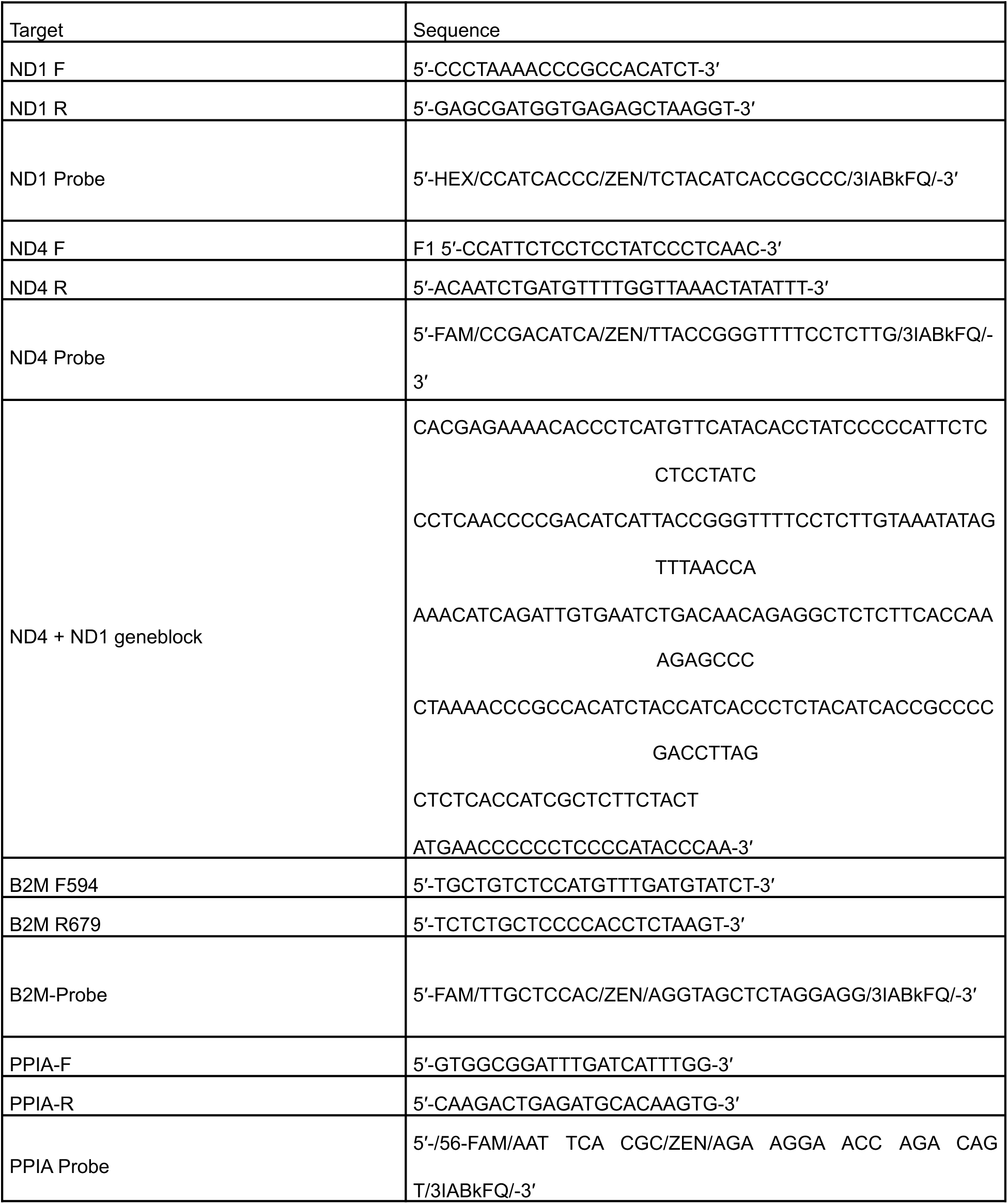
Probes and primer sequences for quantitative PCR detection of ccf mtDNA and mtDNA copynumber. ND1 and ND4 target mitochondrial DNA while B2M and PPIA target nuclear reference genes.

### Electrophysiology Analysis and Calcium Imaging

Neurons were seeded on 48-well black Cytoview MEA plate containing 16 electrodes per well (Axion Biosystems) and electrical activity was measured on the Axion Maestro Pro MEA system (Axion Biosystems). As for calcium imaging, Fluo-4AM (MCE HY-101896) and Rhod-2FM (Invitrogen, R1244) were used for spontaneous calcium recordings. Details on spikes, bursts, and network burst parameters and data acquisition for electrophysiology and calcium are available in Supplemental Methods.

### Statistical Analysis

All statistical analysis methods are listed in each figure and were completed in statistical analysis in GraphPad (Prism 10.6.1) or in Rstudio (Version 2025.05.1+513). All data was assessed for normal distribution and p-value thresholds were set to significance p < 0.05. Principle component analysis (PCA) was conducted on metabolomics, lipidomics, and electrophysiology and linear mixed effect modeling (LME) using estimated marginal means (EMM) was applied to characterize group x stage interactions. More details are described in Supplemental Methods.

## Results

### Clinical, biological, and genetic characteristics of participants

Clinical and demographic information for all participants is summarized in Supplementary Table 1, with CT and BD data adapted from El Sabbagh et al. (2025)^7^. For patients with BD-FMD, details of family relatives who have been clinically diagnosed at the Montreal Children’s Hospital with a mitochondrial disease are listed in Supplementary Figure 1 and Supplementary Table 2.

Whole-genome sequencing (WGS) was performed on all donor-derived iPSC lines to evaluate potential genetic contributors to mitochondrial dysfunction. No shared deleterious variants or pathogenic mtDNA mutations were detected within or across disease groups, indicating that mitochondrial abnormalities in BD and BD-FMD are not homogenously driven by rare coding variants (Supplementary File 1).

### Validation and characterization of iPSC-derived NPCs and CNs across disease groups

IPSC characterization of CT and BD lines are currently published and remaining lines shown in Supplementary Figure 2 and Supplementary Tables 3-4^7^. IPSC-derived NPCs from all groups robustly expressed Nestin (Figure 2B; Supplementary Figure 2D). CNs displayed MAP2⁺ neurite networks with minimal GFAP⁺ astrocytes, and subtype markers identified both VGLUT1⁺ glutamatergic and GABA⁺ GABAergic neurons (Figure 2C-D). Cell-type composition (%VGLUT, %GABA) was comparable across groups (Figure 2E; Supplementary Figures 3-4).

**Figure 2.**
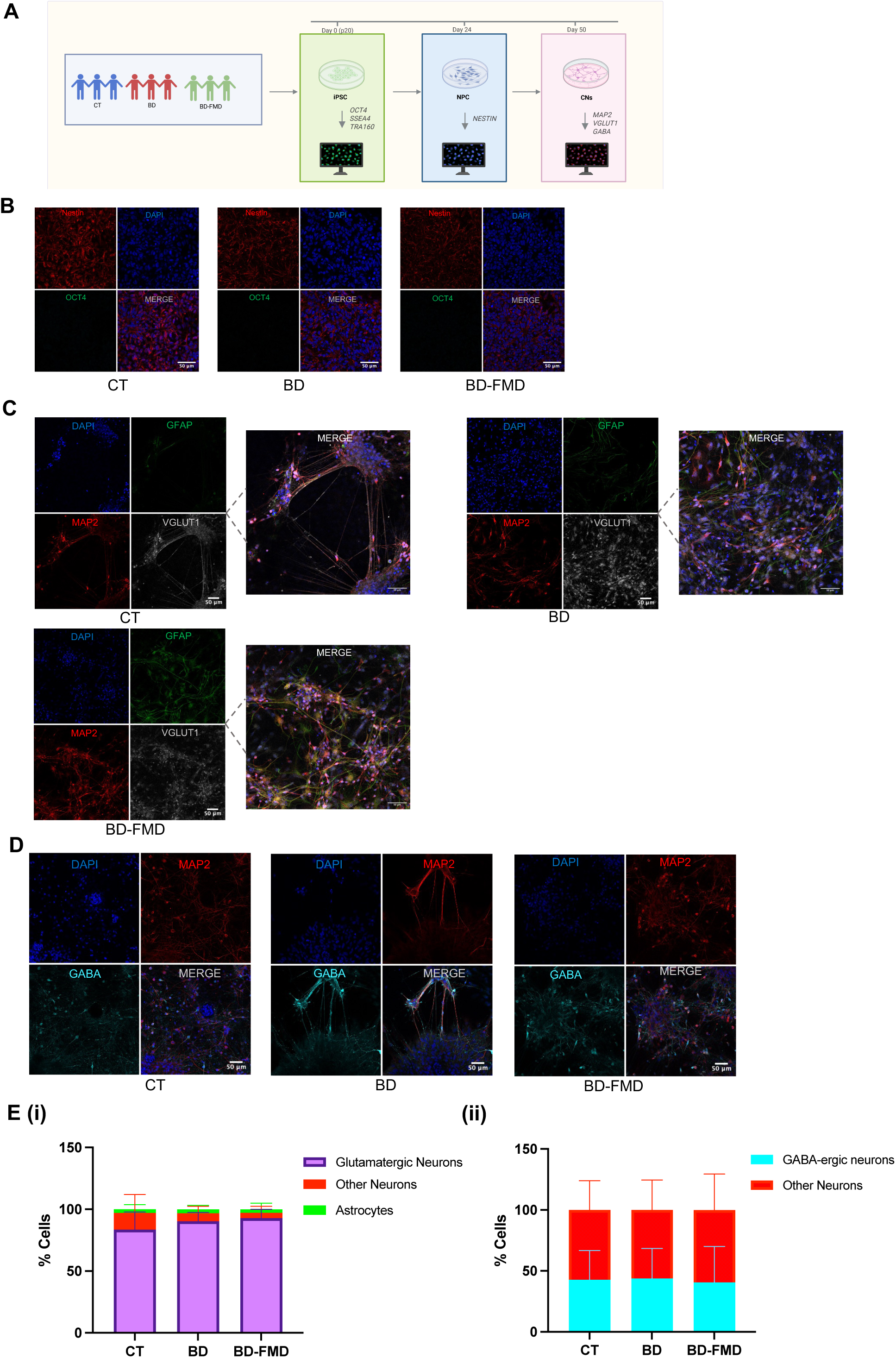
Validation and characterization of iPSC-derived NPCs and CNs across study groups. (A) Schematic of the differentiation workflow from iPSCs to NPCs (day 24; OCT4, NESTIN) and CNs (day 50; MAP2, VGLUT1, GABA, GFAP). (B) Representative immunofluorescence images of NPCs from control (CT), bipolar disorder (BD), and BD with family history of mitochondrial disease (BD-FMD), for OCT4 (green) and Nestin (red) expression (scale bar, 50 μm). (C) Representative immunofluorescence images of cortical glutamatergic neurons identified by MAP2 (red), VGLUT1 (gray), and astrocytic marker GFAP (green) across groups (scale bar, 50μm). (D) Representative immunofluorescence images of GABAergic neurons identified by MAP2 (red) and GABA (cyan) across groups (scale bar, 50μm). (n=3 samples per group x 6 FOV x 3 batches). (E) Quantification of CN subtype composition at day 50. (i) Distribution of glutamatergic neurons (purple), other neurons (red), astrocytes (green), and other unspecified cell types (blue) as percent of total cells (mean ± SD). (ii) Distribution of GABAergic neurons (blue), other neurons (red), and other cell types (gray) as percent of total cells (mean ± SD). Two-way ANOVA with no significant differences reported across groups for proportions of glutamatergic neurons, GABA-ergic neurons or astrocytes.

### Examining Mitochondrial and Cellular Health across iPSCs, NPCs, and CNs

Across groups, six readouts: mitochondrial membrane potential (MMP) (JC-1), ATP production, mtDNA copy number, ROS release, double-stranded DNA (dsDNA) release, and circulating cell-free mtDNA (ccf-mtDNA) (Figures 3B-G, respectively) were assessed across the three cell stages (iPSCs, NPCs, and CNs). In Figure 3, panels (i) in each row show the model-estimated group trajectories from a linear mixed-effects model (LME) and panels (ii) show the actual observed trajectories per group^18^.

**Figure 3.**
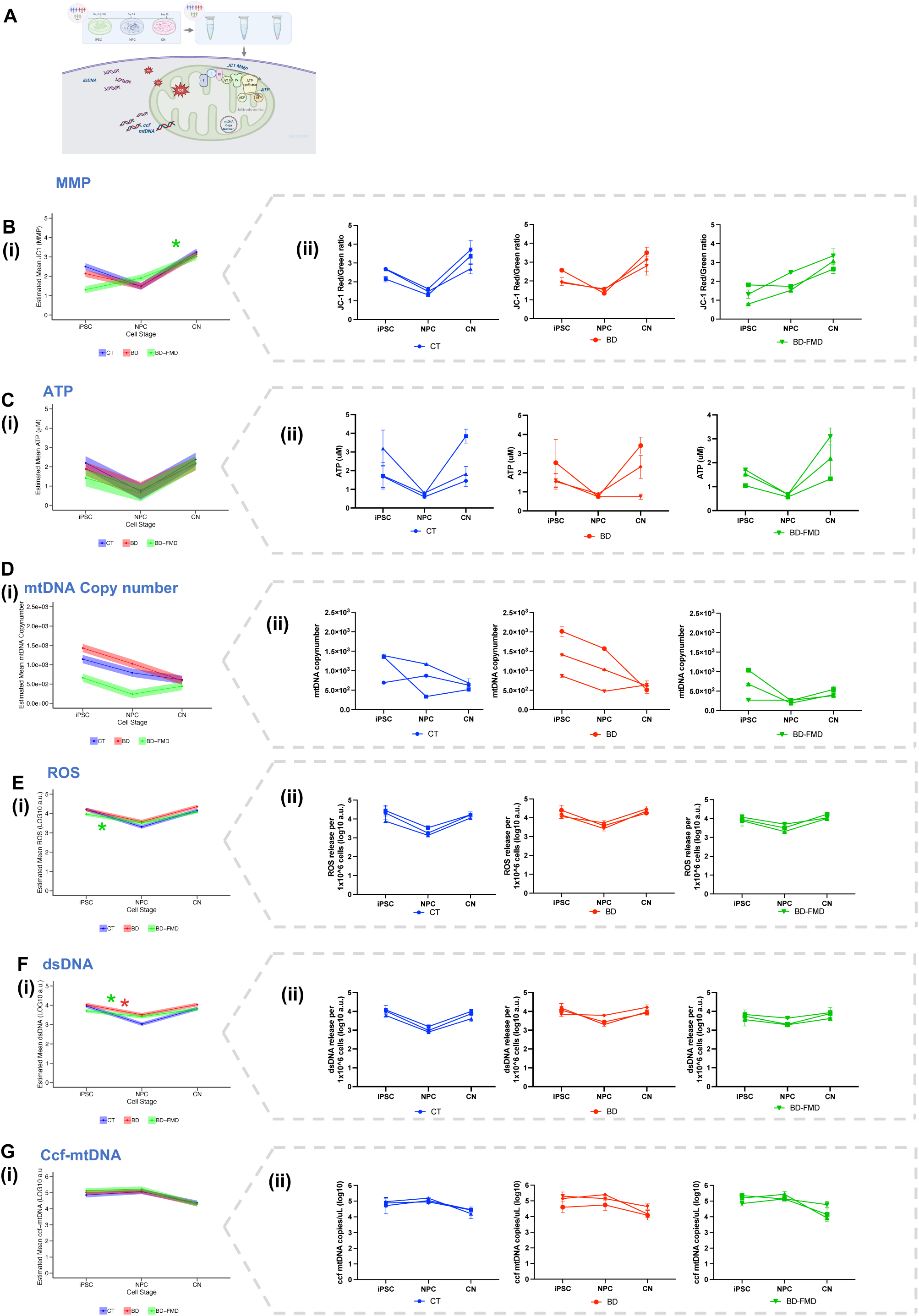
Examining Mitochondrial and Cellular Health across iPSCs, NPCs, and CNs. Schematic overview of measurements done on cell types. (B)-(G) For each assay, the left panel (i) Graphs show fitted value trajectories for CT, BD, BD-FMD obtained from Linear Mixed-Effects Model (LME) including groups and cell stage as factors, and the right panels (ii) Graphs show each group’s observed values at each stage. (B) Mitochondrial membrane potential (MMP) by JC-1 red/green ratio. (ii) GLMM: CT vs BD-FMD in NPC to CN (p=0.005) (C) Cellular ATP concentrations per 20,000 cells. (D) mtDNA copy number values, normalized to nDNA. (E) Relative ROS production. (ii) GLMM: CT vs BD-FMD in iPSC to NPC (p=0.007) (F) Relative extracellular double-stranded DNA release (dsDNA). (ii) GLMM CT vs BD in iPSC to NPC (p=0.02) and CT vs BD-FMD in iPSC to NPC (p=0.0008) (G) Levels of circulating cell free mtDNA (ccf mtDNA) release. Statistics use a LME with CT as a reference group and iPSC as a reference stage.

At the intracellular level, visual inspection, although not significant showed a dip at the NPC stage in the MMP and ATP, and a decline in mtDNA copy number with progression to CNs in all groups (Figures 3Bii, Cii, Dii). Extracellularly, ROS and dsDNA release reached minimum points at the NPC stage, however ccf-mtDNA levels showed broadly parallel patterns across the groups (Figures 3Eii, Fii, Gii).

For MMP in Figure 3Bi, LME identified a significant linear trend group x stage interaction for the NPC to CN stage transition in the BD-FMD group (*p* = 0.005), indicating a magnitude of stage-dependent shift in MMP during NPC to CN differentiation. By contrast, ATP content and mtDNA copy number did not show significant group x stage interactions in BD nor in BD-FMD, suggesting similar stage-dependent patterns across all groups (Figures 3 Ci, Di). In the extracellular environment, ROS release across the iPSC to NPC stage transition was significantly different in BD-FMD (*p* = 0.007) and dsDNA release across the iPSC to NPC stage was significantly altered in BD (*p* = 0.02) and in BD-FMD (*p* = 0.0008), as their trajectories followed a flatter-trend across differentiation compared to CT (Figures 3Ei-Fi). Collectively, BD-FMD showed distinct trajectories in MMP and ROS, and both BD and BD-FMD displayed flatter trends for dsDNA, indicating heightened dsDNA release across neurodifferentiation, while CT’s trend fluctuated based on the cell type.

### Pathway-level analysis identifies distinct metabolomic alterations during NPC to CN differentiation in BD and BD-FMD

Metabolomics data from NPCs and CNs revealed that PC1 (48.2%) primarily captured variance related to cell type, with NPC and CN samples clustering separately regardless of diagnosis. In contrast, PC2 (18.1%) reflected an interaction between diagnosis and differentiation stage (Supplementary Figure 5). While no significant differences were observed between groups at the NPC stage, PC2 scores at the CN stage differed significantly for BD (*p* = 0.0052) and BD-FMD (*p* = 0.047) relative to CT, indicating that disease-specific metabolic alterations emerged during neuronal differentiation. These findings prompted pathway-level over-representation analyses (ORA) to identify the significantly represented metabolic pathways from NPC and CN-specific metabolites during development (Figure 4; Supplementary Figure 6).

**Figure 4.**
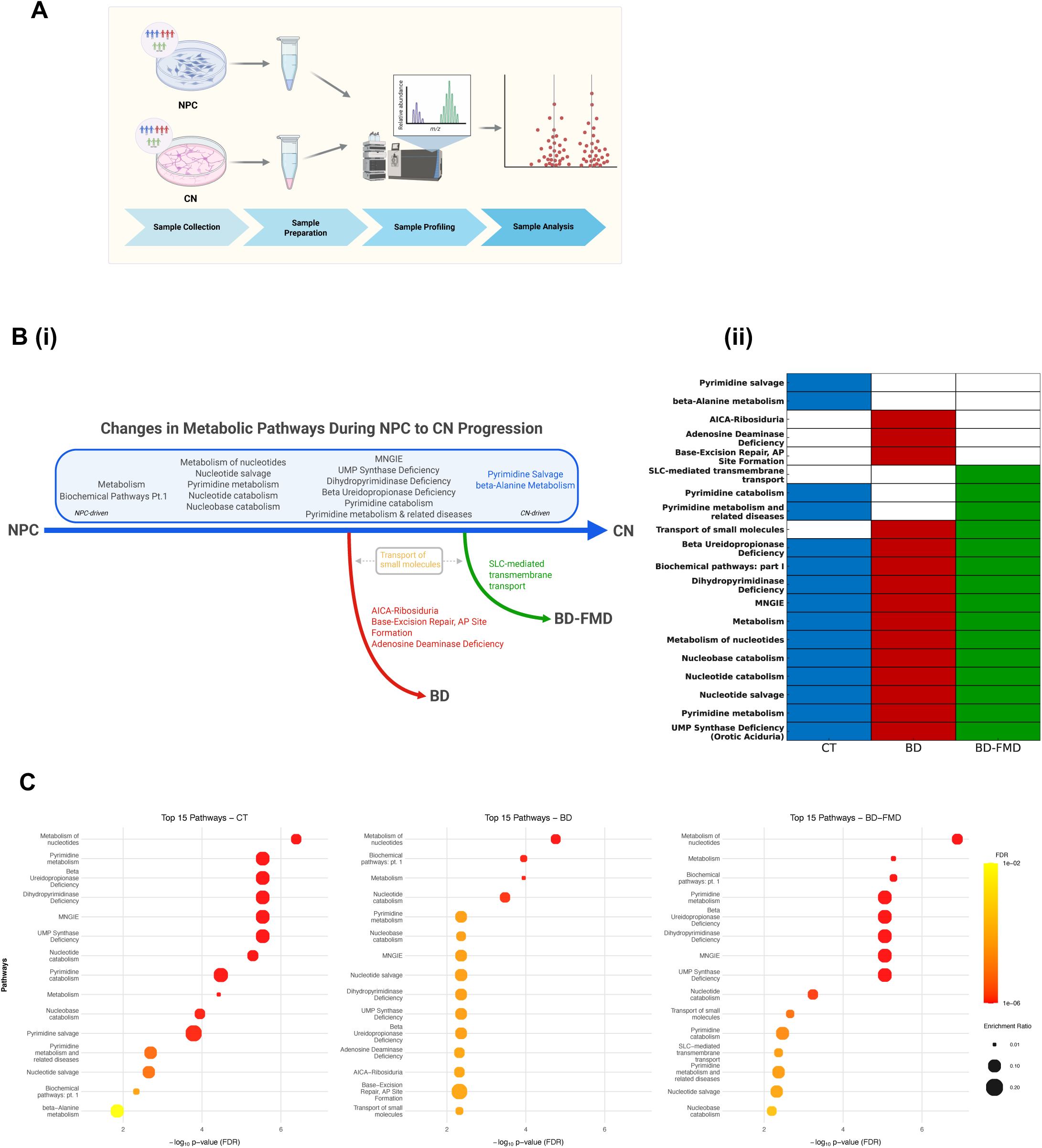
Pathway-level analysis identifies distinct metabolomic alterations during NPC to CN differentiation in BD and BD-FMD. (A) Schematic of the LC/MS workflow. NPC and CN lysates from CT (blue), BD (red), and BD-FMD (green) were profiled. For each group (n=3 donor lines), stage-restricted metabolite sets were defined (present at one stage, absent at the other) and submitted to over-representation analysis (ORA) in MetaboAnalyst (Fisher’s exact test with FDR correction). (B) (i) Summary map of pathway changes across the NPC→CN pipeline, highlighting shared developmental pathways (black), CT-specific pathways (blue), BD-specific deviations (red), and BD-FMD-specific deviations (green), and both disease-shared deviations (yellow). (ii) Presence/absence matrix of enriched pathways by group, illustrating CT-only, BD-only, BD-FMD-only, and shared pathways. (C) Top-15 enriched pathways per group. X-axis: -log10(FDR adjusted p-value). Bubble size = enrichment ratio (hits/expected). Color encodes FDR (red = lower FDR). Abbreviations: FDR, false discovery rate; SLC, solute carrier; AICAR, 5-aminoimidazole-4-carboxamide ribonucleotide. Pathways named after diseases (e.g., ADA deficiency, MNGIE) reflect metabolite sets used for enrichment and are interpreted here as pathway signatures rather than clinical diagnoses.

Eleven pathways were significantly represented across all groups (Figure 4B). Among these, two library-defined categories: Biochemical pathways Part 1 and Metabolism, captured the metabolic remodeling during NPC to CN transition as NPCs require coordinated metabolic rewiring to successfully differentiate into CNs^9^ (*q* < 0.01). Among the shared pathways, pyrimidine core metabolism was a common theme across all groups with defined pathways including pyrimidine metabolism, nucleotide salvage, and metabolism of nucleotides (*q* < 0.01), as these pathways are essential for purine and pyrimidine synthesis, supporting DNA and RNA synthesis, mtDNA maintenance, and neurite outgrowth functions^10,19,20^.

Beyond the shared pathways, group-specific signatures also emerged, driven by the presence or absence of a small set of stage-specific metabolites. Unique pathways that were significantly represented in CT but not in BD or BD-FMD were pyrimidine salvage (*q* < 0.001), supporting DNA and RNA synthesis and β-alanine metabolism (*q* < 0.01), reflecting systematic pyrimidine catabolism^21–23^. Both BD and BD-FMD groups exhibited altered NPC and CN metabolite signatures which suggest altered energy metabolism and impaired neuronal maturation relative to CT. Transport of small molecules (*q* = 0.0048 in BD and *q* = 0.0022 in BD-FMD) was a significantly represented pathway found across both disease groups indicating a hallmark during their neurodifferentiation (Figure 4C).

The BD group also showed a unique subset of significant pathways: adenosine deaminase (ADA) deficiency, base-excision repair, AP site formation, and AICA-ribosiduria (5-aminoimidazole-4-carboxamide) pathways (*q* < 0.01). The BD-FMD group also exhibited a unique subset of NPC-specific metabolites associated with altered activity of the SLC-mediated transmembrane transport pathway (*q* < 0.01). These pathways directly influence substrate levels, proton-ion homeostasis and redox balance^24^. Collectively, the metabolomic profiles define a shared NPC to CN differentiation pattern, with disease-specific deviations in BD and BD-FMD marking distinct, stage-linked trajectories.

Next, to determine whether lipidomic profiles paralleled metabolomic trends, we performed PCA on combined lipidomic data from NPCs and CNs. PC1 (65.15%) primarily separated samples by cell type, reflecting broad lipidomic remodeling during neuronal differentiation. In contrast, PC2 (11.54%) did not distinguish diagnostic groups, and no significant group and cell-type interaction effects were found (Supplementary Figure 7). These results indicate that lipidomic variance is largely driven by developmental stage, with no evidence of disease-related modulation during the NPC-to-CN transition.

### Neurite outgrowth and network MEA electrophysiology diverge across CT, BD, and BD-FMD neurons

To better understand BD and BD-FMD in longitudinal development of neuronal circuits, MEA electrophysiology recordings were used. CNs formed dense, spontaneous active networks across all groups when cultured on MEAs and overall neuronal viability remained similar across groups and across maturation from day 42 to day 70 (Figure 5B).

**Figure 5.**
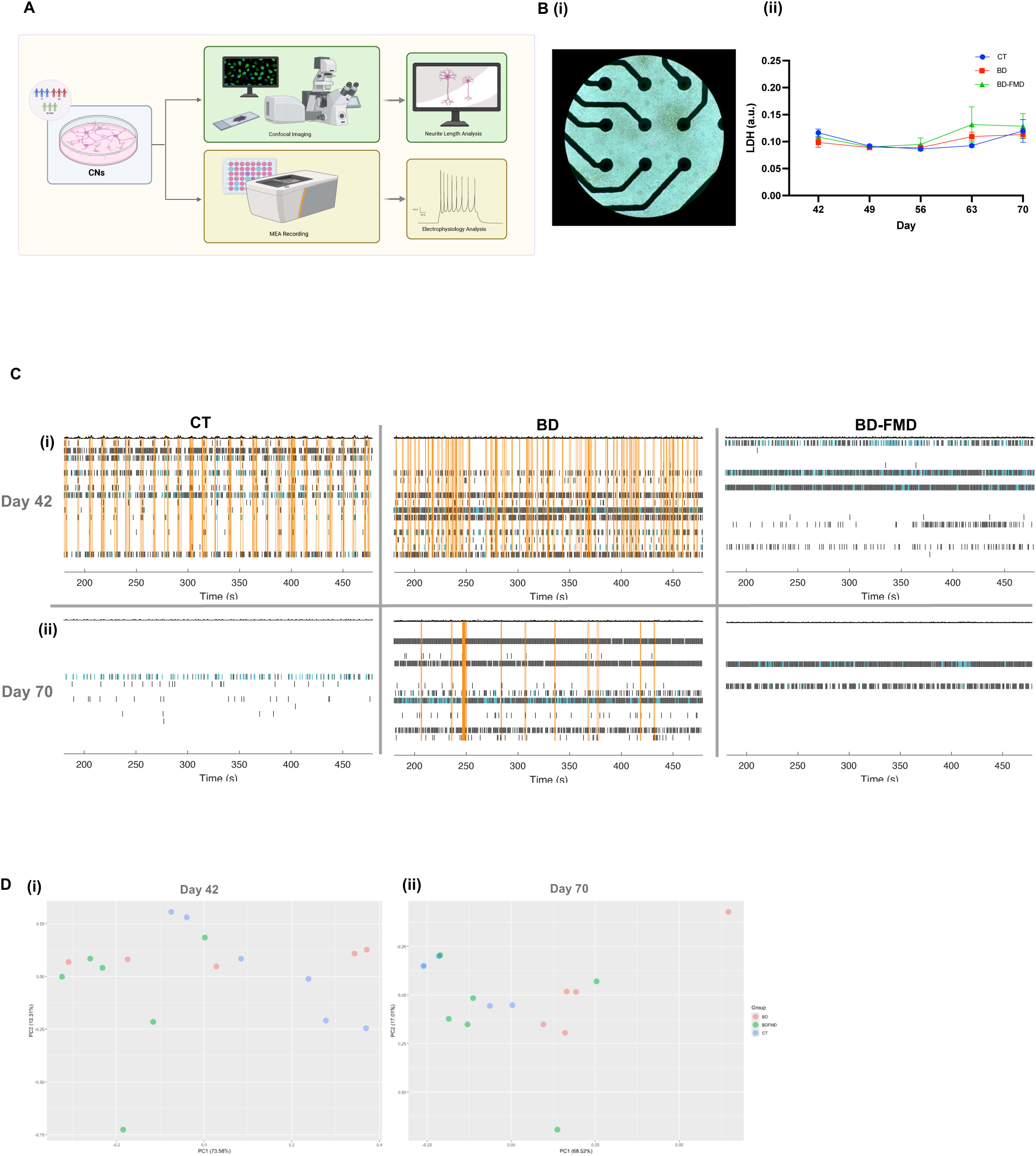
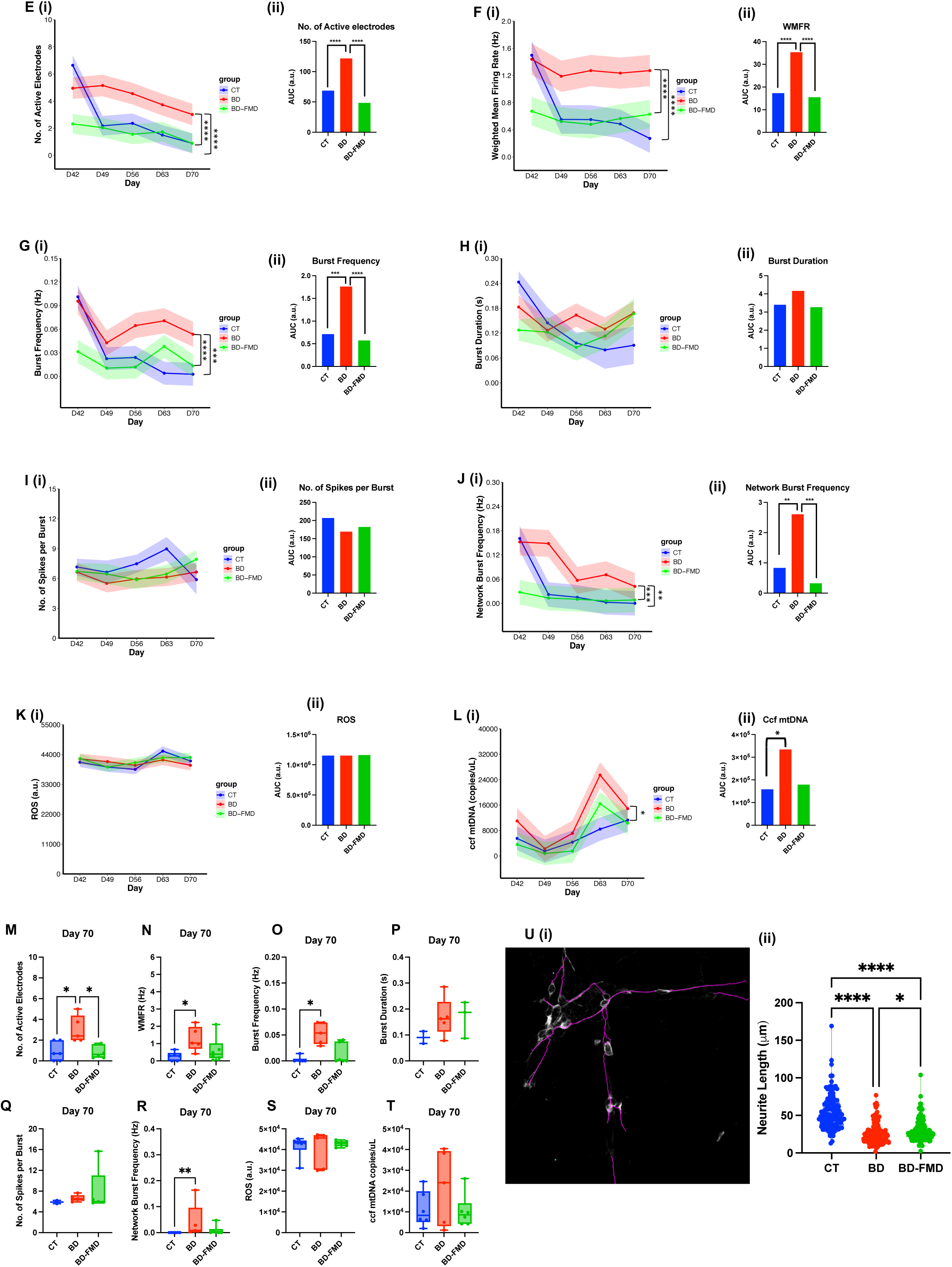
Network MEA electrophysiology activity and neurite outgrowth diverge across CT, BD, and BD-FMD neurons. (A) Schematic of experimental workflow for MEA recordings and neurite length quantification. (B) (i) Representative image of neurons cultured on MEA plate, 10X magnification. (ii) LDH measurements across each recording timepoint. (C) Representative raster plots showing spontaneous spikes, bursts, and network bursts over a 5-minute time segment at Day 42 (i) and Day 70 (ii). (D) Principle component analysis (PCA) of all MEA parameters at Day 42 (i) and Day 70 (ii). (E-L) (i) Linear mixed-effects (LME) analyses of key MEA parameters modeling trajectory from Day 42 to Day 70. (ii) AUC of MEA metric from each corresponding LME plotted. Significance asterisks denote differences in area-under-the-curve (AUC) across groups derived from post-hoc contrasts on the LME model (\**p* < 0.05, \*\**p* < 0.005, \*\**p* < 0.005). (E) number (No.) of active electrodes (CT vs BD *p* < 0.001; BD vs BD-FMD *p* < 0.001), (F) weighted mean firing rate (WMFR) (CT vs BD *p* < 0.001; BD vs BD-FMD *p* < 0.001), (G) burst frequency (CT vs BD *p* < 0.001; BD vs BD-FMD *p* < 0.001), (H) burst duration, (I) spikes per burst, (J) network burst frequency (CT vs BD *p* = 0.007; BD vs BD-FMD *p <* 0.001), (K) reactive oxygen species (ROS) levels, and (L) circulating cell-free mitochondrial DNA (ccf-mtDNA) *(p* = 0.034). Lines represent estimated marginal means (EMM) derived from the LME and shaded areas represent SEM. (M-T) Boxplots representing raw data at Day 70 of each MEA metric. Group differences were analyzed by one-way ANOVA followed by Šidák’s post-hoc correction for multiple comparisons. \**p* < 0.05, \*\**p* < 0.01. N=2 per sample including 4 wells per line, 2 batches per line. Total n=40 wells x 2 batches. (U) (i) representative image of neurite length measurement taken at 20X magnification with SNT Viewer plugin on FIJI. (ii) measurement of neurite length (µm) across disease groups. P<0.0001 for CT vs BD and CT vs BD-FMD; P=0.04 for BD vs BD-FMD. Data analyzed by Kruskal Wallis test and post-hoc Dunn’s multiple comparison test. \**p* < 0.05, \*\**p* < 0.01, \*\*\**p* < 0.005, \*\*\*\**p* < 0.001.

Raster plots of 5-minute recordings revealed activity differences between CT, BD, and BD-FMD where black lines represent spikes, blue lines represent bursts, and orange boxes represent network bursts (Figure 5C). At day 42, CT, BD, and BD-FMD neurons were distinguishable: CT networks separated by both PC1 and PC2, BD networks separated mainly along PC1, BD-FMD mainly along PC2, indicating distinct variance between the three groups. By day 70, this separation changed as BD-FMD samples became more dispersed and partially overlapped with CT, whereas BD remained distinct from both. These temporal shifts suggest that BD and BD-FMD neurons diverge early in network development but follow different maturation trajectories (Figure 5D).

For quantification, the AUC of each LME model’s EMM was computed to determine cumulative electrophysiological activity across development (Fig. 5E-L). The number of active electrodes decreased in all groups over development; however, this reduction was not due to reduced neuronal survival (Figure 5Bii). This late-stage reduction in electrode engagement also aligns with previous studies on iPSC derived neurons cultured on MEA with similar maturation timelines^25^. The magnitude of this decline differed markedly between groups as BD neurons maintained a significantly higher number of active electrodes than both CT and BD-FMD across development (*p* < 0.001) (Figure 5E). Weighted mean firing rate (WMFR) was examined, which adjusts spike counts by the fraction of active electrodes. BD neurons exhibited persistently higher WMFR than both CT and BD-FMD throughout development (*p* < 0.001) (Figure 5F). This aligns with the disease-specific segregation observed in the PCA, reinforcing that BD and BD-FMD neurons maintain different electrophysiological signatures, despite comparable developmental timelines.

Next, AUC of burst frequency followed the same trend as WMFR, remaining consistently higher in BD neurons relative to CT and BD-FMD across time (*p* < 0.001) (Figure 5G). Burst duration and spikes per burst showed no significant group differences in AUC measurements (Figures 5H-I), indicating that although BD neurons fired more frequently, the temporal structure of individual bursts was preserved. Network bursts were also examined since this parameter has shown to be increased in BD-derived GABAergic neurons^26^. Network burst frequency mirrored the burst frequency pattern (Figure 5J), remaining elevated in BD neurons compared to both CT and BD-FMD (*p* = 0.007 and *p* < 0.001, respectively). ROS and ccf-mtDNA were quantified from the supernatant collected directly after MEA recordings. ROS did not differ significantly between groups (Figure 5K), suggesting comparable environments during basal firing conditions, however ccf-mtDNA was elevated in BD (Figure 5L) (*p =* 0.034).

Endpoint measurements at day 70 were also consistent with the AUC-based comparisons (Figures 5M-T). The BD group exhibited higher number of active electrodes and a higher WMFR compared to CT and BD-FMD (*p* < 0.05), consistent with hyperexcitability at later stages (Figures 5M-N). BD also exhibited significantly higher burst frequency and sustained high network bursts compared to CT (*p* < 0.05 and *p* < 0.01) while BD-FMD cultures exhibited intermediate bursts indicating partial normalization of network activity (Figures 5O, R).

Neurite length was also measured and was significantly different across groups. Figure 5U shows neurite lengths of CT was significantly longer than BD-FMD (*p* = 0.04) and BD (*p* < 0.001). BD showed the shortest neurite length overall. Together, these findings indicate both BD and BD-FMD neurons display reduced neuronal complexity and higher excitability patterns compared to CT, however the impact is more pronounced in BD.

### Cytosolic and mitochondrial calcium dynamics reveal disease specific differences across CT, BD, and BD-FMD cortical neurons

To determine whether these network-level differences arise from altered intracellular calcium (Ca²⁺) dynamics and mitochondrial signaling, we next performed simultaneous Ca²⁺ imaging using cytosolic Fluo-4 AM and mitochondrial Rhod-2 AM indicators^27,28^. Mitochondrial health has been known to affect changes in intracellular Ca²⁺, especially in patients with BD^1^. In the CNs, cytosolic Ca²⁺ transients measured using Fluo-4 revealed significance group differences in peak amplitude (Figure 6B). BD-FMD neurons exhibited higher peak amplitudes (denoted as ΔF/F0) compared to both CT and BD neurons, indicating enhanced cytosolic Ca²⁺ responses (*p* = 0.0002 for CT vs BD-FMD; *p* = 0.04 for BD vs BD-FMD) (Figures 6C). Peak frequency and width did not differ significantly across groups (Figures 6D-E). Mitochondrial Ca²⁺ dynamics were also assessed using Rhod-2 (Figure 6F). While overall amplitude and frequency of Ca²⁺ activity were comparable across groups, BD-FMD neurons displayed broader peak widths relative to CT (*p* = 0.0032), indicating prolonged mitochondrial Ca²⁺ retention (Figures 6G-I).

**Figure 6.**
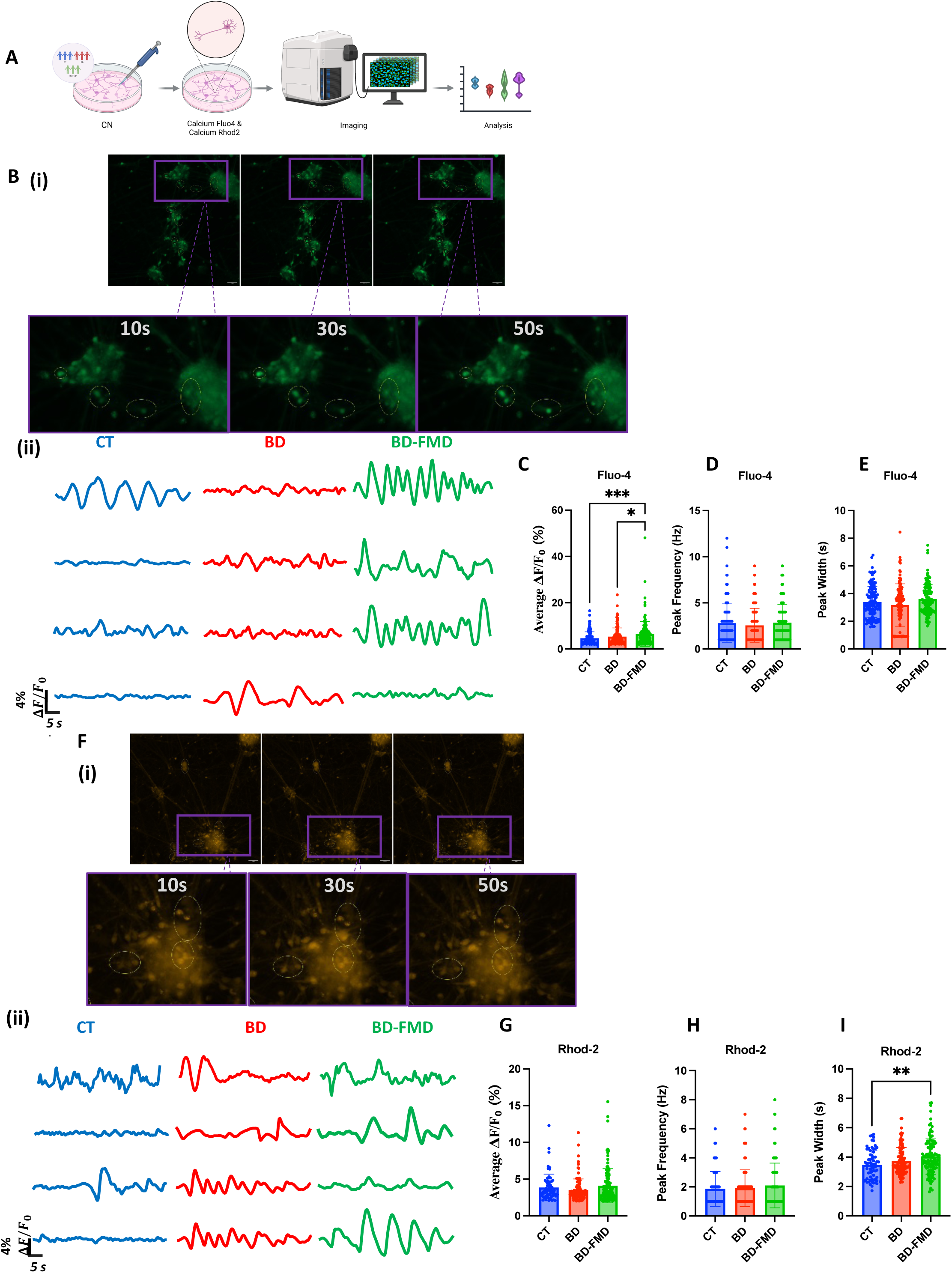
Cytosolic and mitochondrial calcium dynamics reveal disease specific differences across CT, BD, and BD-FMD cortical neurons. (A) Schematic of experimental workflow using Fluo-4 and Rhod-2 calcium indicators. (B) (i) Representative Fluo-4AM fluorescence images showing spontaneous calcium transients in cortical neurons at 10, 30, and 50 seconds (20X magnification) Images cropped for visual representation (Purple lines). (ii) Representative ΔF/F₀ traces from individual cells across groups. (C-E) Quantification of Fluo-4 parameters: (C) average ΔF/F₀ (%) (*p* = 0.0002 for CT vs BD-FMD; *p* = 0.04 for BD vs BD-FMD), (D) peak frequency (Hz), and (E) peak width (s). (F) (i) Representative Rhod-2AM fluorescence images showing spontaneous calcium transients in cortical neurons at 10, 30, and 50 seconds (20X magnification) Images cropped for visual representation (Purple lines). (ii) Representative ΔF/F₀ traces. (G-I) Quantification of Rhod-2 parameters: (G) average ΔF/F₀ (%), (H) peak frequency (Hz), and (I) peak width (s). CT vs BD-FMD *p* = 0.0032. All data analyzed by Kruskal-Wallis test with Dunn’s multiple-comparison post-hoc test. Bar graphs show the mean ± SD.

These experiments reveal distinct functional profiles across CNs derived from CT, BD, and BD-FMD patient lines. While BD neurons exhibit hyperexcitable electrophysiological patterns, BD-FMD neurons displayed reduced overall network activity but increased cytosolic and mitochondrial Ca²⁺ responses. These results highlight the complementary patterns of neuronal dysfunction where one is driven by network hyperactivity and the other by altered Ca²⁺ handling.

## Discussion

BD is currently a top ten cause of global disability with high rates of premature mortality from suicide and medical comorbidities^29^. Patients with BD-FMD represent a mitochondrial-risk enriched stratum of BD, where a familial mitochondrial burden, coming from the same mitochondrial disease (COX deficiency), plausibly adds to a psychiatric vulnerability beyond what is found in BD without FMD. The PMD COX deficiency is a complex IV mitochondrial disorder causing impaired respiration, membrane instability, and increased ROS leaks^30^. Hence, this study delineates risk-context specific differences in mitochondrial susceptibility and stage-resolved metabolomic pathways to identify unique molecular characteristics of BD-FMD in comparison to BD across neurodifferentiation trajectories. Briefly, our results suggest that among patients with BD, familial mitochondrial disease may contribute to a distinct etiopathophysiology through developmental, metabolic, and functional mechanisms.

Within this framework, disease and stage-specific metabolic signatures emerged when differentiating from iPSC to NPC to CNs. LME modeling revealed a stronger dependence on mitochondrial regulation in BD-FMD with different fluctuations in ROS and MMP during differentiation. These changes across neurodifferentiation in BD-FMD are not the same as in the CTs, unmasking the familial mitochondrial liability when OXPHOS demands rise^9^. Altered ROS release can also cause ROS-induced bursts which then affect dsDNA release, both of which were detected during the iPSC-to-NPC transition in BD-FMD. These alterations suggest that mitochondrial stress phenotypes are detectable even when clinical penetrance is incomplete. Consistent with PMD literature showing elevated ROS, impaired respiration and altered MMP, the BD-FMD lines represent a mitochondrial-fragility phenotype that can intersect with psychiatric vulnerability^12,31–34^. However, the dsDNA alteration in the BD group may reflect both a lighter mitochondrial dysfunction load, coupled with inflammation-linked mechanisms rather than overt mitochondrial instability^7,35,36^.

Molecular alterations can also lead to excessive proliferation of NPCs or premature differentiation to CNs which can give rise to neurodevelopmental disorders^37^. Metabolic profiles change within cell stages and these can influence cell fates and functions^10^. Therefore, examining key significantly represented metabolic pathways during the NPC to CN stage progression is crucial to understand stage and disease specific differences. Metabolomic ORA primarily captured a stage driven program where most significant pathways overlapped across groups and were tied to nucleotide metabolism (*de novo* and salvage synthesis), consistent with NPC to CN transition’s need to balance DNA/RNA biosynthesis^10^. While this ORA identified significant differences across groups, the directionality of these changes cannot be inferred from ORA alone. These findings reflect that specific metabolic pathways are perturbed across diagnostic groups, while accounting for stage-specific changes that normally occur in healthy neurodifferentiation.

Disease-specific pathways emerged against this developmental baseline. BD and BD-FMD both had alterations implicated in the transport of small molecule, which regulate metabolite exchange across cell membranes^38^. This finding was notable given reports linking BD and PMD to abnormal transporter and neurotransmitter regulation^39^. The BD-specific involvement in ADA-linked pathways and base-excision repair points to purine turnover and DNA repair activity. ADA converts adenosine to inosine, regulating intracellular ATP levels^3,40^. Altered ADA activity has been previously reported in bipolar mania and purine metabolism abnormalities associated with pharmacotherapy resistance in mood disorders^3,40^. Patients with BD and their relatives have shown increased abnormalities in base-excision repair pathways, consistent with greater genotoxic stress at the NPC stage^40^. Overall, the evidence supports a neurodifferentiation (NPC to CN) predisposition of nuclear DNA damage as reflected in both LME analyses of dsDNA and ORA of metabolomic pathways.

The BD-FMD-specific involvement centered on SLC-mediated transmembrane transport. In this pathway the SLC25 mitochondrial carrier gene family is present with over 50 inner mitochondrial membrane transporters^41^. This pathway includes uncoupling proteins (UCP1-5) and SLC25(A7/8/9/27) which regulate substrate exchange, a proton motive force (MMP), ATP supply, and mitochondrial biogenesis^24,41,42^. Thus, this metabolic pathway signature of BD-FMD provides a biochemical context for the group-stage interaction seen in BD-FMD (Figure 3B and 3E) on the MMP and ROS levels during iPSC to NPC to CN differentiation as SLC activity regulates substrate supply, ion/proton homeostasis, inner mitochondrial membrane proteins, ketone levels like acetoacetate, and AMPK signaling^24,43^. Together, these results support a model in which familial mitochondrial risk interacts with the metabolic demands of neurodifferentiation (BD-FMD), while inflammatory/repair pathways contribute a partly distinct signature in BD detectable as changes in extracellular dsDNA levels.

Extending beyond pathway-level analysis, structural and functional readouts were consistent with these metabolic interpretations. Mitochondrial dysfunction and PMD have been previously shown to drive neuronal dysregulation phenotypes, and this may explain why BD-FMD CNs showed a low activity profile across development^44^. BD-FMD CNs fit an energy-limited model of reduced mitochondrial support, and diminishing axonal and synaptic upkeep. BD CNs, however, have a different set of unique metabolites and exhibit a hyperexcitable state at later maturation stages, consistent with studies of hyperexcitability in BD iPSC derived neurons and COs^7,26,45–47^.

The differences in electrophysiological trajectories of BD and BD-FMD may also reflect the distinct underlying mechanisms of Ca²⁺ regulation. Although BD neurons uniquely exhibit hyperexcitability and higher network activity, their cytosolic and mitochondrial Ca²⁺ amplitudes remain relatively stable. Compensatory buffering mechanisms, such as enhanced Ca²⁺ extrusion or rapid mitochondrial Ca²⁺ cycling, may be limiting large Ca²⁺ spikes during individual firing events^48^. However, this compensation likely comes at a metabolic cost: the cumulative, activity-driven Ca²⁺ load, which over time may impose chronic mitochondrial stress. This interpretation is supported with the elevated ccf-mtDNA release observed between days 42 and 70 in BD, reflecting a secondary, activity-induced mitochondrial stress rather than an intrinsic mitochondrial defect. In contrast, BD-FMD neurons display exaggerated cytosolic Ca²⁺ peaks and prolonged mitochondrial Ca²⁺ retention despite lower network activity, indicating intrinsic Ca²⁺ handling dysfunction rather than activity-dependent stress. These abnormalities are not uncommon as there are known mitochondrial vulnerabilities which have previously shown to impair the ER-mitochondria communication complex between the voltage-dependent anion channel (VDAC), the inositol 1,4,5-trophosphate receptor (IP₃R), and molecular chaperone glucose-regulated protein 75 (GRP75)^49^. Such disturbances weaken the coordination of Ca²⁺ exchange required for ATP production and redox stability, potentially predisposing neurons to energy failure and oxidative imbalance^49^.

Previous studies have similarly shown that Ca²⁺ handling in BD neurons varies by developmental stage, with neural progenitors exhibiting smaller Ca²⁺ transients and mature neurons displaying exaggerated spontaneous activity, possibly reflecting maturation-dependent shifts in mitochondrial buffering capacities^6,50^. Sustained Ca²⁺ dysregulation has well-established consequences for neuronal health^51^. Elevated intracellular Ca²⁺can activate both intrinsic and extrinsic apoptotic pathways that eliminate neurons unable to maintain homeostasis^52^. Excess Ca²⁺ perturbs not only excitability but also Ca²⁺-dependent signaling cascades controlling dendritic growth, synaptic plasticity, and excitation-inhibition balance^53^. Together, these findings point to two mechanistic subtypes of neuronal vulnerability in bipolar neurons: a hyperactive, metabolically strained network phenotype (BD) and a hypoactive, calcium dysregulated phenotype (BD-FMD).

## Limitations

This study provides a multi-level characterization of mitochondrial and neuronal function across BD, BD-FMD, and control iPSC-derived lineages, but several limitations should be noted. The sample size was small, reflecting the challenge of obtaining well-characterized BD with familial mitochondrial disease, and thus limits this study. Analyses were based on group-level trajectories due to a small sample size for inferential statistics, therefore replication across larger cohorts will be essential. While the use of iPSC-derived neuronal models has helped to uncover BD and mitochondrial specific phenotypes, the models can only capture trait biology, not manic or depressive episodes. They also are 2D cell models lacking vasculature and peripheral immune, metabolic, and endocrine tissue. MEA activity and calcium transients in iPSC-derived cortical neurons capture early network bursts, not adult oscillations, and the lack of long-range connectivity and myelination limits the capability of these 2D neuronal models. Finally, another limitation is the lack of iPSC-derived cortical neurons from patients with mitochondrial disease only which may have offered a unique perspective of mitochondrial disease-specific neuronal traits.

## Conclusions

Our findings reveal that familial mitochondrial liability actively reshapes neuronal differentiation and function, producing distinct trajectories of mitochondrial performance, calcium signaling, and network excitability. These results suggest that mitochondrial dysfunction in BD is not a secondary byproduct of illness, but a mechanistic contributor to altered neuronal energetics, communication and symptom expression.

Importantly, this work provides a potential translational framework for patient stratification in BD. The observation that calcium signaling and mitochondrial metabolism are jointly perturbed raises the possibility that existing mood stabilizers with mitochondrial or calcium-modulating actions, such as valproic acid, lithium, or nimodipine, may exert differential efficacy in patients with underlying mitochondrial vulnerability (e.g., BD-FMD). Future studies integrating pharmacological perturbations in iPSC-derived neurons could identify biomarker signatures predicting drug responsiveness and guide personalized treatment approaches.

Looking ahead, combining this iPSC-based platform with multi-omic profiling and live-cell bioenergetic assays will enable discovery of molecular nodes linking mitochondrial regulation to synaptic activity. Ultimately, mapping how mitochondrial and calcium networks interact across neurodevelopment could advance precision medicine in psychiatry, transforming the way bipolar disorder is classified and treated.

## Acknowledgements

We would like to thank Alaa Shamandy for coding assistance in python, Rstudio, and MATLAB, and Pavel Powlowski for help with cortical neuron maintenance. All figures were created in BioRender with license: Zachos, K. (2026) https://BioRender.com/0d2w2b0.

## Funding

This research was supported by grants from the Canadian Institutes of Health Research (CIHR #505547 and Brain and Behavior Research Foundation (NARSAD Independent Investigator Award) to A.C.A. and MITO2i Graduate Scholarship to D.E.S.E.S.; M.B. is supported by a NHMRC Leadership 3 Investigator grant (GNT2017131).

## Disclosures

All authors have no disclosures.

## Notes

### Competing Interest Statement

The authors have declared no competing interest.

## References

1 Young, A. H. & Juruena, M. F. The Neurobiology of Bipolar Disorder. Current Topics in Behavioral Neurosciences (2020). 10.1007/7854_2020_179

2 AA, N., et al. Diagnosis and Treatment of Bipolar Disorder: A Review - PubMed. JAMA 330 (10/10/2023). 10.1001/jama.2023.18588

3 Daniels, S. D. & Boison, D. Bipolar Mania and Epilepsy Pathophysiology and Treatment May Converge in Purine Metabolism: A New Perspective on Available Evidence. Neuropharmacology 241 (2023 Oct 9). 10.1016/j.neuropharm.2023.109756

4 Aghababaie-Babaki, P. et al. Global, regional, and national burden and quality of care index (QCI) of bipolar disorder: A systematic analysis of the Global Burden of Disease Study 1990 to 2019. International Journal of Social Psychiatry 69 (2023-06-23). 10.1177/00207640231182358

5 Phalnikar, K. et al. Altered neuroepithelial morphogenesis and migration defects in iPSC-derived cerebral organoids and 2D neural stem cells in familial bipolar disorder. Oxford Open Neuroscience 3 (2024/02/01). 10.1093/oons/kvae007

6 Hewitt, T. et al. Bipolar disorder-iPSC derived neural progenitor cells exhibit dysregulation of store-operated Ca2+ entry and accelerated differentiation. Molecular Psychiatry 2023 28:12 28 (2023-07-04). 10.1038/s41380-023-02152-6

7. El Soufi El Sabbagh, D. et al. iPSC-derived cerebral organoids reveal mitochondrial, inflammatory and neuronal vulnerabilities in bipolar disorder. Translational Psychiatry 2025 15:1 15 (2025-08-25). 10.1038/s41398-025-03529-7

8 Kathuria, A. et al. Transcriptome analysis and functional characterization of cerebral organoids in bipolar disorder. Genome Medicine 2020 12:1 12 (2020-04-19). 10.1186/s13073-020-00733-6

9 Garone, C. et al. Mitochondrial metabolism in neural stem cells and implications for neurodevelopmental and neurodegenerative diseases. Journal of Translational Medicine 2024 22:1 22 (2024-03-04). 10.1186/s12967-024-05041-w

10 Scandella, V., Petrelli, F., Moore, D. L., Braun, S. M. G. & Knobloch, M. Neural stem cell metabolism revisited: a critical role for mitochondria. Trends in Endocrinology & Metabolism 34 (2023/08/01). 10.1016/j.tem.2023.05.008

11 Bernard, J. et al. Mitochondria at the crossroad of dysregulated inflammatory and metabolic processes in bipolar disorders. Brain, Behavior, and Immunity 123 (2025/01/01). 10.1016/j.bbi.2024.10.008

12 Inczedy-Farkas, G. et al. Psychiatric symptoms of patients with primary mitochondrial DNA disorders. Behavioral and Brain Functions : BBF 8 (2012 Feb 13). 10.1186/1744-9081-8-9

13 Kasahara, T. & Kato, T. What Can Mitochondrial DNA Analysis Tell Us About Mood Disorders? Biological Psychiatry 83 (2018/05/01). 10.1016/j.biopsych.2017.09.010

14 Rosella, L. C. et al. A population-based cohort study of mitochondrial disease and mental health conditions in Ontario, Canada. Orphanet Journal of Rare Diseases 20 (2025 Apr 14). 10.1186/s13023-025-03688-2

15 Colasanti, A. et al. Primary mitochondrial diseases increase susceptibility to bipolar affective disorder. Journal of Neurology, Neurosurgery & Psychiatry 91 (2020-08-01). 10.1136/jnnp-2020-323632

16 IL, K., et al. Cognitive functioning and mental health in mitochondrial disease: A systematic scoping review - PubMed. Neuroscience and biobehavioral reviews 125 (2021 Jun). 10.1016/j.neubiorev.2021.02.004

17 Kato, T. Neurobiological basis of bipolar disorder: Mitochondrial dysfunction hypothesis and beyond. Schizophrenia Research 187 (2017/09/01). 10.1016/j.schres.2016.10.037

18 Bates, D., Mächler, M., Bolker, B. & Walker, S. Fitting Linear Mixed-Effects Models Using lme4. Journal of Statistical Software 67 (2015/10/07). 10.18637/jss.v067.i01

19 Yan, J., Wu, J., Xu, M., Wang, M. & Guo, W. Disrupted de novo pyrimidine biosynthesis impairs adult hippocampal neurogenesis and cognition in pyridoxine-dependent epilepsy. Science Advances 10 (2024-04-05). 10.1126/sciadv.adl2764

20 Wanrooij, P. H. & Chabes, A. NME6: ribonucleotide salvage sustains mitochondrial transcription. The EMBO Journal 42 (2023-08-07). 10.15252/embj.2023114990

21 Bröer, S. & Gether, U. The solute carrier 6 family of transporters. British Journal of Pharmacology 167 (2012 Sep). 10.1111/j.1476-5381.2012.01975.x

22 Mori, M., Gähwiler, B. H. & Gerber, U. β-Alanine and taurine as endogenous agonists at glycine receptors in rat hippocampus in vitro. The Journal of Physiology 539 (2002 Feb 15). 10.1113/jphysiol.2001.013147

23 GH, M. & GP, J. Biochemical Mechanisms of Beneficial Effects of Beta-Alanine Supplements on Cognition - PubMed. Biochemistry. Biokhimiia 88 (2023 Aug). 10.1134/S0006297923080114

24 Schumann, T. et al. Solute Carrier Transporters as Potential Targets for the Treatment of Metabolic Disease. Pharmacological Reviews 72 (2020/01/01). 10.1124/pr.118.015735

25 Brown, C. O. et al. Disruption of the autism-associated gene SCN2A alters synaptic development and neuronal signaling in patient iPSC-glutamatergic neurons. Frontiers in Cellular Neuroscience 17 (2024 Jan 16). 10.3389/fncel.2023.1239069

26 DJ, S., et al. Human-Induced Pluripotent Stem Cell (iPSC)-Derived GABAergic Neuron Differentiation in Bipolar Disorder - PubMed. Cells 13 (07/15/2024). 10.3390/cells13141194

27 KR, G., et al. Chemical and physiological characterization of fluo-4 Ca(2+)-indicator dyes - PubMed. Cell calcium 27 (2000 Feb). 10.1054/ceca.1999.0095

28 Paredes, R. M., Etzler, J. C., Watts, L. T. & Lechleiter, J. D. Chemical Calcium Indicators. Methods (San Diego, Calif.) 46 (2008 Oct 16). 10.1016/j.ymeth.2008.09.025

29 Robinson, N. et al. Impact of Early-Life Factors on Risk for Schizophrenia and Bipolar Disorder. Schizophrenia Bulletin 49 (2023 Mar 22). 10.1093/schbul/sbac205

30 Garone, C. Mitochondrial Cytochrome C deficiency can show the first disease signs in the prenatal stage. European Journal of Human Genetics 31 (2023 Oct 4). 10.1038/s41431-023-01466-x

31 RE, A., SL, G., MA, T., MF, M. & PI, R. The psychiatric manifestations of mitochondrial disorders: a case and review of the literature - PubMed. The Journal of clinical psychiatry 73 (2012 Apr). 10.4088/JCP.11r07237

32 Andreazza, A. C., Shao, L., Wang, J.-F. & Young, L. T. Mitochondrial Complex I Activity and Oxidative Damage to Mitochondrial Proteins in the Prefrontal Cortex of Patients With Bipolar Disorder. Archives of General Psychiatry 67 (2010/04/01). 10.1001/archgenpsychiatry.2010.22

33 Leipnitz, G. et al. Evaluation of mitochondrial bioenergetics, dynamics, endoplasmic reticulum-mitochondria crosstalk, and reactive oxygen species in fibroblasts from patients with complex I deficiency. Scientific Reports 2018 8:1 8 (2018-01-18). 10.1038/s41598-018-19543-3

34 Distelmaier, F. et al. The antioxidant Trolox restores mitochondrial membrane potential and Ca2+-stimulated ATP production in human complex I deficiency. Journal of Molecular Medicine 2009 87:5 87 (2009-03-03). 10.1007/s00109-009-0452-5

35 Shabab, T., Khanabdali, R., Moghadamtousi, S. Z., Kadir, H. A. & Mohan, G. Neuroinflammation pathways: a general review. Int J Neurosci 127, 624–633 (2017). 10.1080/00207454.2016.1212854

36 Kim, H. K., Andreazza, A. C., Elmi, N., Chen, W. & Young, L. T. Nod-like receptor pyrin containing 3 (NLRP3) in the post-mortem frontal cortex from patients with bipolar disorder: A potential mediator between mitochondria and immune-activation. J Psychiatr Res 72, 43–50 (2016). 10.1016/j.jpsychires.2015.10.015

37 E, K., F, F., D, J. & S, C. Mapping the molecular and cellular complexity of cortical malformations - PubMed. Science (New York, N.Y.) 371 (01/22/2021). 10.1126/science.aba4517

38 S, H., R, H., T, L., M, R. & P, S. A dopamine transporter mutation associated with bipolar affective disorder causes inhibition of transporter cell surface expression - PubMed. Molecular psychiatry 10 (2005 Dec). 10.1038/sj.mp.4001730

39 Beccano-Kelly, D. A. et al. Calcium dysregulation combined with mitochondrial failure and electrophysiological maturity converge in Parkinson’s iPSC-dopamine neurons. iScience 26 (2023/07/21). 10.1016/j.isci.2023.107044

40 Arat Çelik, H. E., et al. Oxidatively-induced DNA base damage and base excision repair abnormalities in siblings of individuals with bipolar disorder DNA damage and repair in bipolar disorder. Translational Psychiatry 2024 14:1 14 (2024-05-24). 10.1038/s41398-024-02901-3

41 Ogunbona, O. B. & Claypool, S. M. Emerging Roles in the Biogenesis of Cytochrome c Oxidase for Members of the Mitochondrial Carrier Family. Frontiers in Cell and Developmental Biology 7 (2019 Jan 31). 10.3389/fcell.2019.00003

42 Palmieri, F. Mitochondrial transporters of the SLC25 family and associated diseases: a review. Journal of Inherited Metabolic Disease 37 (2014/07/01). 10.1007/s10545-014-9708-5

43 A, K., et al. Akt and AMPK activators rescue hyperexcitability in neurons from patients with bipolar disorder - PubMed. EBioMedicine 104 (2024 Jun). 10.1016/j.ebiom.2024.105161

44 Denley, M. C. S. et al. Mitochondrial dysfunction drives a neuronal exhaustion phenotype in methylmalonic aciduria. Communications Biology 2025 8:1 8 (2025-03-11). 10.1038/s42003-025-07828-z

45 J, M., et al. Differential responses to lithium in hyperexcitable neurons from patients with bipolar disorder - PubMed. Nature 527 (11/05/2015). 10.1038/nature15526

46 S, S., et al. Neurons derived from patients with bipolar disorder divide into intrinsically different sub-populations of neurons, predicting the patients’ responsiveness to lithium - PubMed. Molecular psychiatry 23 (2018 Jun). 10.1038/mp.2016.260

47 R, S., et al. Deficient LEF1 expression is associated with lithium resistance and hyperexcitability in neurons derived from bipolar disorder patients - PubMed. Molecular psychiatry 26 (2021 Jun). 10.1038/s41380-020-00981-3

48 Lee, S. H., Duron, H. E. & Chaudhuri, D. Beyond the TCA cycle: new insights into mitochondrial calcium regulation of oxidative phosphorylation. Biochemical Society transactions 51 (2023 Aug 31). 10.1042/BST20230012

49 Patergnani, S. et al. Calcium signaling around Mitochondria Associated Membranes (MAMs). Cell Communication and Signaling : CCS 9 (2011 Sep 22). 10.1186/1478-811X-9-19

50 McCarthy, M. J. et al. Calcium Channel Genes Associated with Bipolar Disorder Modulate Lithium’s Amplification of Circadian Rhythms. Neuropharmacology 101 (2015 Oct 22). 10.1016/j.neuropharm.2015.10.017

51 Giménez-Palomo, A. et al. Mitochondrial Dysfunction as a Biomarker of Illness State in Bipolar Disorder: A Critical Review. Brain Sciences 14 (2024 Nov 28). 10.3390/brainsci14121199

52. Sousa, R. T. d., Machado-Vieira, R., Carlos A Zarate, J. & Manji, H. K. Targeting mitochondrially mediated plasticity to develop improved therapeutics for bipolar disorder. Expert opinion on therapeutic targets 18 (2014 Jul 24). 10.1517/14728222.2014.940893

53. K, P., I, D., M, M. & N, T. Mitophagy and age-related pathologies: Development of new therapeutics by targeting mitochondrial turnover - PubMed. Pharmacology & therapeutics 178 (2017 Oct). 10.1016/j.pharmthera.2017.04.005

