## Supplemental Methods for "From iPSCs to NPCs to cortical neurons: bioenergetic, neuronal, and calcium signaling phenotypes in bipolar disorder with and without familial mitochondrial disease"

El Soufi El Sabbagh *et al.*

**Methods and Materials**

Study Design

IPSCs were generated from the 9 subjects and then differentiated into NPCs and CNs, enabling stage-specific disease type comparisons. Mitochondrial health was profiles across stages using mitochondrial membrane potential assessments, ATP production, ROS production, and extracellular mtDNA release. In NPCs and CNs, stage-specific metabolomics were examined along with ORA of metabolic pathways. Neuronal identity and composition were quantified with functional readouts including micro-electrode array (MEA) electrophysiology network activity and calcium imaging dynamics.

Blood Sample Processing and iPSC Generation

Peripheral blood mononuclear cells (PBMCs) were isolated from blood collected and reprogrammed into iPSCs using episomal vector reprogramming as described by Duong et al (2021)^1^. All involved participants provided written consent to participate in this study (University of Toronto REB #29949). All details on blood samples and processing information for CT and BD group has been published by El Sabbagh et al (2025) and followed the same methods and reprogramming steps for BD-FMD iPSC lines^2^. Characterization of iPSCs was done by karyotyping and examining the protein expression of pluripotent markers (SSEA4, OCT4, SOX2, TRA-1-60) using the Pluripotent Stem Cell 4-Marker Immunocytochemistry Kit (Invitrogen, A24881) for immunofluorescence. This study was approved by the Research Ethics Board at the University of Toronto, Ontario, Canada using REB (36359) and REB (29949) and at Deakin University with REB (17/205) in accordance with the Helsinki Declaration of 1975. Dr. Andreazza’s laboratory has also received approval from the Stem Cell Oversight Committee (Canadian Institutes of Health Research, #399222).

Whole Genome Sequencing (WGS)

Whole genome sequencing of all donor samples was performed at the Next Generation Sequencing Facility at The Centre for Applied Genomics (TCAG, SickKids, Toronto, Canada). Qubit dsDNA HS Assay was used to quantify genomic DNA and sample purity was assessed through a OD260/OD280 ratio. Illumina TruSeq PCR-free DNA library (Illumina) was used with 700 ng of DNA following the manufacturer’s protocol. A Bioanalyzer High Sensitivity DNA Chip was used for assessing final libraries and this was quantified by qPCR by the KAPA Library Quantification Illumina/ACI Prism protocol from Kapa Biosystems. these validated libraries were sequenced in one lane per sample following Illumina’s recommended protocol on a HiSeq X.

Data processing of WGS data was performed by the Bioinformatics facility at TCAG where the software bcl2fastq, provided by Illumina, was used to convert BCL base call files from the Illumina Sequencing systems to a standard primary sequencing in 38 FASTQ format and these metrics were computed to examine overall quality of experiments. All reads were aligned to the reference human genome UCSC hg19 by the Isaac Aligner (iSAACSAAC00776.15.01.27). Nuclear-encoded mitochondrial genes were filtered using MitoCarta 3.0, and variants were annotated based on established pathogenicity thresholds (pLI > 0.9, CADD_phred > 30, PolyPhen > 0.9)^3-5^.

Karyotyping of iPSCs

Karyotype was done as previously described in Duong et al. (2021)^1^. In brief, karyotype analysis was performed by G-banding on iPSCs expanded in two 25-cm² flasks and processed by the TCAG Cytogenomics Facility. At ~50% confluence, cells were treated with Karyomax Colcemid (0.15 µg/mL) for 30 or 60 minutes, dissociated with trypsin–EDTA, and subjected to hypotonic treatment (0.054 M KCl, 20 mM HEPES, 0.02% EGTA) at 37 °C for 25 minutes. Cells were then fixed in Carnoy’s fixative (methanol/acetic acid, 3:1) with multiple fixation rounds, dropped onto glass slides, and aged prior to G-banding. Twenty metaphase spreads per line were evaluated following standard cytogenetic procedures.

Generating NPCs from iPSCs

NPC were generated from iPSCs using an in-house protocol. First, **N2 medium** was prepared by combining DMEM/F-12 with GlutaMAX™ and phenol red (Invitrogen; 97.25%), N-2 Supplement (1X; 1:100 dilution) (Thermo Fisher 17502048), GlutaMAX™ Supplement (1 mM) (Thermo Fisher 35050061), MEM Non-Essential Amino Acids (100 μM) (Thermo Fisher 111140050), and 2-mercaptoethanol (0.1 mM; 1:500 dilution) (Thermo Fisher 21985023). Second**, B27 medium** was prepared using Neurobasal™ Plus medium (98%) (Thermo Fisher A3582901), supplemented with B-27™ Supplement without vitamin A (1X; 1:50 dilution) (Thermo Fisher 125870) and GlutaMAX™ (1 mM; 1:200 dilution). Finally, **Neural Induction Medium** was generated by mixing equal parts of N2 and B27 media (50:50). The medium was further supplemented with the following small molecules: **LDN193189** (100 nM) (Tocris 6053), **SB431542** (10 µM) (Tocris 1614), and **XAV939** (2 µM) (Tocris 3748). All small molecules were added fresh daily to maintain activity and media was changed every day for 10 days. Cells were replated on day 11 and from days 10-16, small molecules added included **LDN193189 (**100 nM), **SB431542** (10 µM), human recombinant bFGF (10 µg/mL) (StemCell technologies 78003) and human recombinant EGF (10 µg/mL) (StemCell technologies 78006). Media was changed daily and on day 16 cells were re-plated with Neural induction medium containing only human recombinant bFGF (10 µg/mL) and human recombinant EGF (10 µg/mL). Daily media changes were done with fresh additions of small molecules and growth factors and by day 21 they were used for experiments.

Protein Expression

To validate iPSC and NPC identity at the mRNA level, we performed quantitative reverse transcription PCR (qRT-PCR). Pre-validated RT² qPCR primer assays (Qiagen) targeting SOX2, POUF5F1, KLF4, MYCL1, LIN28A were used for iPSCs and pre-validated RT² qPCR primer assays (Qiagen) targeting **NES, SOX2, PAX6, and POUF5F1**, along with the reference genes **GAPDH** and **ACTB**, were used for NPCs. Total RNA was isolated from iPSCs and NPCs using the RNeasy Mini Kit (Qiagen), and cDNA was generated from 500 ng of RNA using the RT² First Strand Kit according to the manufacturer’s instructions. Each reaction contained 1 µL of cDNA, 1 µL of the respective RT² primer assay, 12.5 µL of RT² SYBR Green Master Mix, and 10.5 µL of nuclease-free water. Amplification was carried out on a Bio-Rad CFX96 system using the following cycling conditions: 95 °C for 10 min (enzyme activation), followed by 45 cycles of 95 °C for 15 s and 60 °C for 1 min for fluorescence acquisition. Gene expression was normalized to the geometric mean of **GAPDH** and **ACTB** and reported as **2^⁻ΔCt^,** where

ΔCt = Ct(target gene) - Ct[GEOMEAN(GAPDH, ACTB)].

Generating cortical neurons

Cortical neurons were generated using the Forebrain Neuron Differentiation Kit (StemCell Technologies 08600). When NPCs were 21 days old, they were seeded at a density of 2 million cells per mL and differentiated to neural precursors for 7 days. Neural precursors were then passaged once again and seeded for maturation at a density of 3 million cells per mL using the Forebrain Neuron Maturation Kit (StemCell Technologies 08605). Cortical neurons matured up to day 70 with half media changes occurring every 2 days.

Immunofluorescence

IPSCs, NPCs, and CNs were fixed with 4% paraformaldehyde (PFA) (8 minutes for iPSC and NPC, 15 min for CNs) and then washed three times with PBS, 5 minutes each. Samples were permeabilized and blocked with 3% bovine serum albumin (BSA) and 0.1% Triton-X. Antibodies were diluted in 0.5% BSA and after staining coverslips were mounted on glass slides using ProLong Gold Antifade Mountant containing DAPI (Invitrogen, Waltham, MA, USA, P36935). A complete list of antibodies used in this study are provided in Table 1.

Confocal imaging and image analysis

Immunofluorescent imaging was performed at the Krembil Brain Institute, Toronto, Canada using an LSM 880 Elyra Super resolution Confocal Microscope. Neurons were generated in 3 batches and in each batch, 6 fields of view were taken to ensure maximal representation of cells. Confocal images were acquired under identical acquisition settings across samples. Each image was subsequently analyzed using **HALO** (Indica Labs, v4.0.5107.357) using the **Cytonuclear FL module** (v4.2.3), which enables automated detection, segmentation, and quantification of individual cells based on nuclear and cytoplasmic fluorescence. Two independent staining panels were analyzed: (1) **VGLUT1/MAP2/GFAP,** and (2) **GABA/MAP2** to identify inhibitory neurons.

Singularizing cells for downstream analyses

IPSCs grew in colonies and werre singularized using Gentle Cell Dissociation Reagent (GCDR) (StemCell Technologies 100-0485). In brief, iPSCs cultured in 6 well plates were washed twice with PBS, then 1 mL of GCDR was added and cells were put to incubate at 37°C 5% CO_2_ for 8 minutes. After 8 minutes cells were briefly tapped and mixed with media and proceeded to count the singularized iPSCs.

NPCs and CNs which were also grown in 6 well plates followed a similar protocol to the above however Accutase (StemCell Technologies 0792) was used instead of GCDR. Accutase incubation lasted approximately 7 minutes for NPCs and 20 minutes for CNs and cells proceeded for counting.

Intracellular ATP Measurements

Intracellular ATP was measured using Cell Titer Glo Luminescent Assay (Promega G7570) using manufacturer’s protocols. In brief, iPSCs, NPCs, and CNs were seeded at a density of 20,000 cells per well in 100 μL Hanks’ Balanced Salt Solution (HBSS) in a 96 well white plate (Greiner CELLSTAR, 655083). More details on this are available in El Sabbagh et al. (2025)^2^.

JC1 kinetic

iPSCs, NPCs and CNs were dissociated and stained with JC-1 ((5,5′,6,6′-tetrachloro-1, 1′,3,3′-tetraethylbenzimi- dazolylcarbocyanine iodide) Dye (Invitrogen, Mitochondrial Membrane Potential Probe, T3168) at a concentration of 1 μg/mL for 30 min in the dark on an orbital shaker at 37 °C with 5% CO_2_ and humidity. Samples were washed three times with PBS and 25,000 cells per well were plated in a 96 well plate to be measured using a Synergy H1 microplate reader equipped with Gen 5 software (BioTek Instruments, Inc., 253147).

MtDNA Copy Number

A protocol was adapted from Picard et al. for measuring relative mtDNA copy number by calculating the ratio of MT-ND1 gene copy to two copies of β-2 microglobulin^6^. A list of primers and probes is available in Table 2 and more details on the protocol is published in Duong et al^1^.

Quantification of Ccf-mtDNA copies

Cell supernatant (100 μL) was collected, and DNA was extracted using QiaAMP DNA mini kit (Qiagen). The ccf-mtDNA was quantified using Taqman^TM^ Duplex polymerase chain reaction (PCR) with primers and probe targeting β2 M and PPIA for nuclear DNA, and ND1 and ND4 for mitochondrial DNA (Table 2). More details on this experimental protocol are available in El Sabbagh et al (2025)^2^.

Quantification of reactive oxygen species levels and double stranded DNA

Levels of ROS production ere measured using 2′,7′-dichlorodihydrofuorescein diacetate (DCFH-DA) (Sigma-AldrichD6883) from cell culture supernatant. Extracellular release of dsDNA was measured using DNA PicoGreen fluorescent probe (Quant-iT^TM^ PicoGreen^TM^, Thermo Fisher-[P11495](https://www.ncbi.nlm.nih.gov/protein/P11495)). Both assays are described in El Sabbagh et al. (2025)^2^. Fluorescence intensity for both assays was measured using a Synergy H1 microplate reader equipped with Gen 5 software (BioTek Instruments, Inc., Winooski, VT, USA, 253147).

Metabolomic and lipidomic profiling

Metabolomic profiling on NPCs and CNs was conducted in collaboration with the University of Ottawa’s Metabolomics Core Facility using liquid chromatography and mass spectrometry (LC/MS). NPC and CNs were singularized and pelleted to 5 million cells per pellet, and 226 small molecule metabolites were quantified. All data was normalized to metabolite quantity per 1 million cells. Further details on metabolite processing can be found in Zachos et al. (2024)^7^. Quantitative values were used to determine presence or absence within a stage and to compile input lists for ORA.

Within each group, a metabolite was classified as NPC specific if detected in 3/3 donor lines NPC and 0/3 donor lines of CN of the same group, and vice versa for CN-specific metabolites per group. Metabolites not meeting these criteria were excluded from stage analyses. The NPC-specific and CN-specific metabolite lists were combined to form each group’s unique input set for over-representation analysis (ORA) of pathway enrichment where driver metabolites can explain group uniqueness, reported in results (Figure 4 and Supplementary Figure 3). Pathway ORA was performed in MetaboAnalyst using Fisher’s exact test with Benjamini-Hochberg FDR correction against small-molecule libraries SMPDB^8^. Significance was defined as FDR-adjusted *p*-values where *q* < 0.05. For visualization, bubble plots displayed the top 15 pathways per group were selected by ascending FDR with two measures reported: FDR adjusted *p*-value, and enrichment ratio (number of input metabolite matched to pathways over expected metabolites). Lipidomic were also measured by LC/MS. NPC and CNs were singularized and pelleted to 5 million cells per pellet, and 112 lipids were quantified. All data was normalized to metabolite quantity per 1 million cells and followed by PCA for data analysis.

Neurite length measurement

Neurite length was quantified using **SNT (Simple Neurite Tracer) in FIJI**. For each biological sample, multiple fields imaged from immunofluorescence-labeled (MAP2) neuronal cultures. Neurites were semi-automatically traced from the soma to terminal tips with SNT and manually curated to resolve crossings/branch points; ambiguous overlaps and out-of-focus processes were excluded. The **total path length per neuron** was exported in µm and aggregated per sample^9^. Data was plotted and statistical analysis was performed in GraphPad (Prism 10.6.1).

Multi-electrode array (MEA)

Wells were coated with filter sterilized 0.1% Poly(ethyleneimine) (PEI) solution in borate buffer (pH 8.4) (Boric acid Sigma B6768 and PEI Sigma P3143) for 2 hours at room temperature. Wells were then washed four times with water, and the plate was left to dry overnight in the biosafety cabinet. The next day, neurons were seeded at a density of 500,000 neurons per well in 200 μL volume of a 48-well black Cytoview MEA plate containing 16 electrodes per well (Cat# M768-tMEA-48B Axion Biosystems). After 2 hours of seeding cells, an additional 100μL was added to each well to achieve a total volume of 300μL per well. STEMdiff forebrain maturation medium (StemCell 08605) was used with supplementation of filter-sterilized with 7.5mM of D-glucose (Sigma G8270) to achieve a final concentration of 10mM. Half media changes were done every two days and cells were incubated in 37°C at 5% CO_2_.

Electrical activity was recorded once a week, for five weeks using the Axion Maestro Pro MEA system (Axion Biosystems). Day of recording would be the same day of a media change but recording occurred prior to the media change. MEA plates were taken out of the incubator and put inside the Axion Maestro Pro MEA system to equilibrate in an environment of 37°C and 5% CO_2_ for 5 minutes. Recording began and lasted 10 minutes where real-time extracellular spontaneous activity was measured and later use for data processing. A threshold of greater than 5.5 standard deviations was used for spike detection and noise separation, and only active electrodes were used in data analysis since they needed to have a minimum of 5 spikes per minute as previously done by other groups^10^. Wells that had no active electrodes for 3 consecutive weeks were excluded from analysis. Following previously published MEA analysis protocols, bursts were defined as a minimum of 5 spikes pin a maximum 100 millisecond inter-spike interval (ISI) and network bursts were defined as a minimum of 10 spikes in a maximum of 100 millisecond period with at least of 35% electrodes in synchrony^10^. Spike data was analyzed with the Axis Navigator software (Axion Biosystems) at a sampling 12.5 kHz with a 4 kHz Kaiser Window low pass filter and a 200 Hz IIR High Pass filter^11^. Analyzed data were exported to CSV files and processing was done using Axion Biosystems Neural Metric Tool and Axis Plotting Tool. Ccf-mtDNA and levels of ROS were measured form cell culture supernatant of the MEA wells following methods described above.

Lactate dehydrogenase (LDH) assay

LDH was measured using cell supernatant from MEA wells and the LDH Cytotoxicity Assay Kit (Cayman Chemicals 601170) following manufacturer’s protocol. Absorbances were measured at a wavelength of 490 nm using a Synergy H1 microplate reader equipped with Gen 5 software (BioTek Instruments, Inc., 253147).

Calcium imaging

Cortical neurons were seeded at a density of 1.5x10^6^ cells/well in Poly-L-ornithine (Sigma P4957) and laminin (Sigma L2020) coated imaging dishes (Ibidi,81156-400) in a media volume of 1.5 mL per well. At Day 50, cells were washed once in warm HBSS (Gibco 14025092), followed by incubation with 1mL HBSS containing 2 μM cytoplasmic Ca²⁺ dye, Fluo-4 AM (MCE HY-101896), and 3 μM mitochondrial Ca²⁺ dye Rhod-2 AM (Invitrogen, R1244). Incubation was for 30 min at 37°C, 5% CO_2_. Cells were washed three times with warm HBSS, then 1mL of HBSS was added to wells. Imaging dishes were imaged using a Zeiss AxioObserver 7 microscope (Zeiss, Germany) equipped with a live imaging chamber at 37°C, 5% CO_2_. Images taken at 3 frames per second (3 FPS) following previous work on iPSC derived neurons in mood disorders where a 1 minute recording was enough to determine baseline spontaneous Ca²⁺ transients for neurons ^12,13^.

Ca²⁺ data analysis was conducted using a custom MATLAB (R2025a Update 1v 29.1.0.2973110 64-bit maca64) code modified from previously published protocols^14-16^. In brief, time-lapse Ca²⁺ imaging videos were first motion-corrected using the NoRMCorre algorithm implemented in MATLAB to eliminate lateral drift and frame misalignment^16^. After motion correction, regions of interest (ROIs) corresponding to individual neuronal somata were manually defined based on the maximum intensity projection of each field on FIJI/ ImageJ^17^. For each ROI, fluorescence intensity traces were extracted and normalized to the baseline fluorescence (F₀), calculated as the mean of the first 20 frames. Changes in fluorescence were calculated as ΔF/F₀ = (F − F₀)/F₀. Peak detection was performed on each ROI’s ΔF/F₀ trace using the MATLAB ‘findpeaks’ function. The frequency of Ca²⁺ transients per cell, as well as their amplitude (peak height) and duration (full width at half maximum denoting peak width) were subsequently calculated. These neuronal measurements were exported for downstream statistical analysis to GraphPad (Prism 10.6.1). The MATLAB Code is available in Supplementary File 2. Absolute quantification of intracellular and mitochondrial Ca²⁺ was also not possible because both Fluo-4 AM and Rhod-2 AM are non-ratiometric dyes. The fluorescence signals therefore represent normalized ΔF/F₀ traces that indicate relative temporal dynamics of Ca²⁺ transients within the cytosol and mitochondria.

Statistical Analysis

Mitochondrial Markers

LME models were generated via *R package lme4^18^* on 3 stages (iPSC, NPC and CNs) with 3 groups (CT, BD, BD-FMD). LME was performed on six markers: MMP, ROS, ATP, mtDNA copy number, dsDNA, and ccf-mtDNA using the reference at the CT iPSC and any interactions were tested to find group and stage specific differences, relative to the reference. To characterize the factor-wise interactions, post-hoc analysis was calculated using estimated marginal means (EMM) in an R package *(Estimated Marginal Means)^19^.*

MEA results

LME models were also used for each MEA metric to assess longitudinal changes across time and between groups. The model included Group (CT, BD, BD-FMD), Time (D42 to D70) and their interaction (Group × Time) as fixed effects. Model fits were used to estimate group-level trajectories and perform pairwise contrasts at each time point. To quantify overall temporal activity, the area under the curve (AUC) for each fitted LME EMM was calculated per group and per metric using trapezoidal method, providing a single integrated measure of electrophysiological maturation over time.

Metabolomics and lipidomics

Missing values from the metabolomics dataset were imputed using half of the threshold for the limit of detection (LOD) or minimum-detected value as provided by manufacturer. Principal component analysis (PCA) was conducted separately on the combined metabolomics and lipidomic dataset after z-transformation. PCA was performed using the prcomp() function in R, and PC1 and PC2 were retained for visualization. For visualization, samples were grouped by disease status (CT, BD, BD-FMD) and cell type (NPC, CN). PC scores were plotted with condition on the x-axis and PC value on the y-axis, with group means connected across NPC and CN stages to illustrate developmental trajectories. PC1 accounted for the majority of variance and aligned with the transition from NPC to CN, whereas PC2 captured subtler, group-specific variance, especially prominent in the CN stage.

PCA on MEA

PCA was also performed on MEA metrics to capture the multivariate electrophysiological profile of each line at early (D42) and later (D70) stages of network development. Metrics included WMFR, burst frequency, network burst frequency, burst duration, spikes per burst, and number of active electrodes. Prior to PCA, all variables were mean-centered and scaled to unit variance. PCA was conducted separately for each time point to reduce dimensionality and visualize group clustering based on overall activity patterns. Principal component loadings were used to identify which electrophysiological parameters contributed most to the separation of groups, and score plots were generated to compare CT, BD, and BD-FMD trajectories at network onset (D42) versus maturation (D70).

**Supplemental References**

1	Duong, A. *et al.* Characterization of mitochondrial health from human peripheral blood mononuclear cells to cerebral organoids derived from induced pluripotent stem cells. *Sci Rep* **11**, 4523 (2021). <https://doi.org/10.1038/s41598-021-84071-6>

2	El Soufi El Sabbagh, D. *et al.* iPSC-derived cerebral organoids reveal mitochondrial, inflammatory and neuronal vulnerabilities in bipolar disorder. *Translational Psychiatry 2025 15:1* **15** (2025-08-25). <https://doi.org/10.1038/s41398-025-03529-7>

3	Lek, M. *et al.* Analysis of protein-coding genetic variation in 60,706 humans. *Nature 2016 536:7616* **536** (2016-08-17). <https://doi.org/10.1038/nature19057>

4	Wang, H. *et al.* Whole-genome sequencing analysis reveals new susceptibility loci and structural variants associated with progressive supranuclear palsy. *Molecular Neurodegeneration* **19** (2024 Aug 16). <https://doi.org/10.1186/s13024-024-00747-3>

5	Rentzsch, P., Witten, D., Cooper, G. M., Shendure, J. & Kircher, M. CADD: predicting the deleteriousness of variants throughout the human genome. *Nucleic Acids Research* **47** (2019/01/08). <https://doi.org/10.1093/nar/gky1016>

6	Picard, M. *et al.* Mitochondrial functions modulate neuroendocrine, metabolic, inflammatory, and transcriptional responses to acute psychological stress. *Proceedings of the National Academy of Sciences* **112** (2015-12-1). <https://doi.org/10.1073/pnas.1515733112>

7	Zachos, K. A. *et al.* Mitochondrial Biomarkers and Metabolic Syndrome in Bipolar Disorder. *Psychiatry Research* **339** (2024/09/01). <https://doi.org/10.1016/j.psychres.2024.116063>

8	Pang, Z. *et al.* MetaboAnalyst 6.0: towards a unified platform for metabolomics data processing, analysis and interpretation. *Nucleic Acids Research* **52**, W398-W406 (2024). <https://doi.org/10.1093/nar/gkae253>

9	Arshadi, C. *et al.* SNT: a unifying toolbox for quantification of neuronal anatomy. *Nature Methods 2021 18:4* **18** (2021-04-01). <https://doi.org/10.1038/s41592-021-01105-7>

10	Brown, C. O. *et al.* Disruption of the autism-associated gene SCN2A alters synaptic development and neuronal signaling in patient iPSC-glutamatergic neurons. *Frontiers in Cellular Neuroscience* **17** (2024 Jan 16). <https://doi.org/10.3389/fncel.2023.1239069>

11	Unda, B. K. *et al.* Impaired OTUD7A-dependent Ankyrin regulation mediates neuronal dysfunction in mouse and human models of the 15q13.3 microdeletion syndrome. *Molecular Psychiatry* **28**, 1747-1769 (2023). <https://doi.org/10.1038/s41380-022-01937-5>

12	Parnell, E. *et al.* Excitatory Dysfunction Drives Network and Calcium Handling Deficits in 16p11.2 Duplication Schizophrenia Induced Pluripotent Stem Cell–Derived Neurons. *Biological Psychiatry* **94** (2023/07/15). <https://doi.org/10.1016/j.biopsych.2022.11.005>

13	Xiong, C. *et al.* Human Induced Pluripotent Stem Cell Derived Sensory Neurons are Sensitive to the Neurotoxic Effects of Paclitaxel. *Clinical and Translational Science* **14** (2021/03/01). <https://doi.org/10.1111/cts.12912>

14	Balachandar, L. *et al.* Simultaneous Ca2+ imaging and optogenetic stimulation of cortical astrocytes in adult murine brain slices. *Current protocols in neuroscience* **94** (2020 Dec). <https://doi.org/10.1002/cpns.110>

15	L, B., C, M., A, S. & JR, D. Characterization of Optimal Optogenetic Stimulation Paradigms to Evoke Calcium Events in Cortical Astrocytes - PubMed. *eNeuro* **12** (09/15/2025). <https://doi.org/10.1523/ENEURO.0220-25.2025>

16	EA, P. & A, G. NoRMCorre: An online algorithm for piecewise rigid motion correction of calcium imaging data - PubMed. *Journal of neuroscience methods* **291** (11/01/2017). <https://doi.org/10.1016/j.jneumeth.2017.07.031>

17	Omelchenko, A. A. *et al.* TACI: an ImageJ Plugin for 3D Calcium Imaging Analysis. *Journal of visualized experiments : JoVE* (2022 Dec 16). <https://doi.org/10.3791/64953>

18	Bates, D., Mächler, M., Bolker, B. & Walker, S. Fitting Linear Mixed-Effects Models Using lme4. *Journal of Statistical Software* **67**, 1 - 48 (2015). <https://doi.org/10.18637/jss.v067.i01>

19	*Estimated Marginal Means: GitHub*, <<https://rvlenth.github.io/emmeans/>> (
