## Supplementary material for "From iPSCs to NPCs to cortical neurons: bioenergetic, neuronal, and calcium signaling phenotypes in bipolar disorder with and without familial mitochondrial disease": Supplemenrary Figures and Tables

### Supplementary Figures

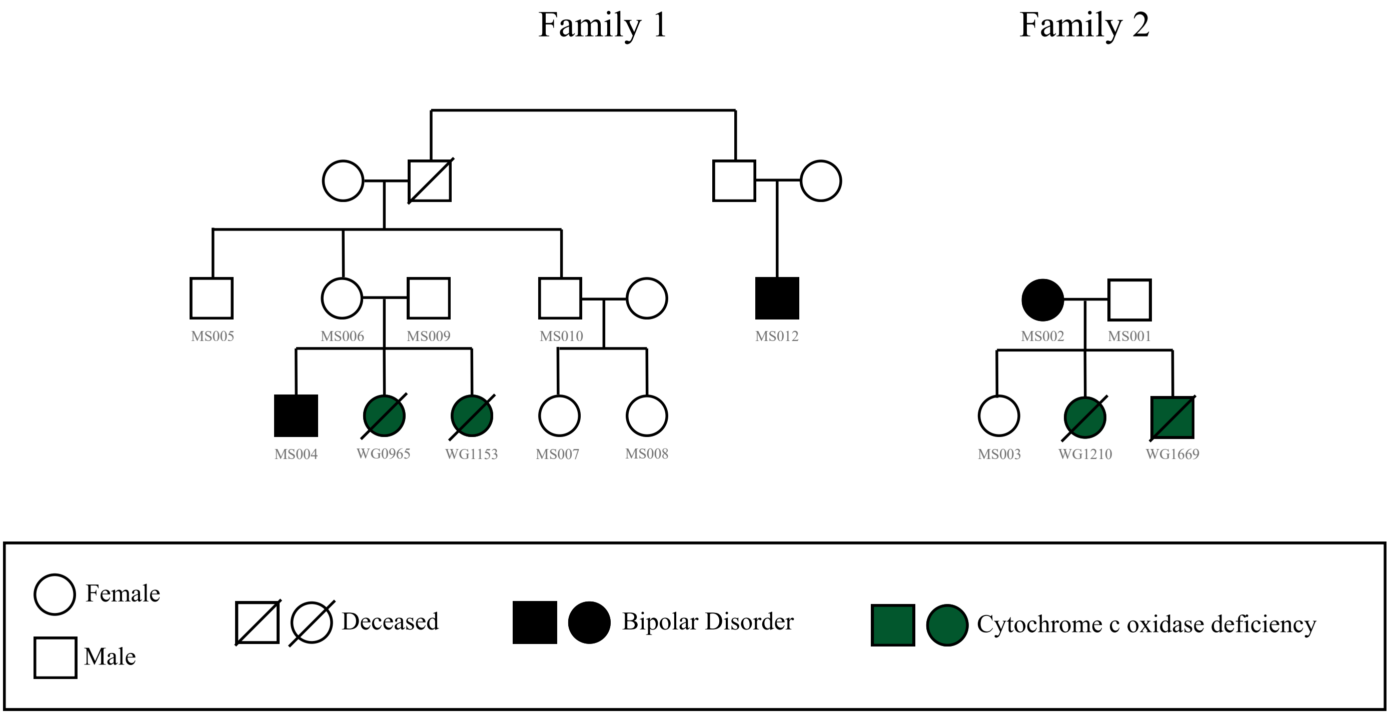


**Supplementary Figure 1.** Pedigree of families. All subjects with informed consent who donated samples were given a sample ID code represented below the symbols.

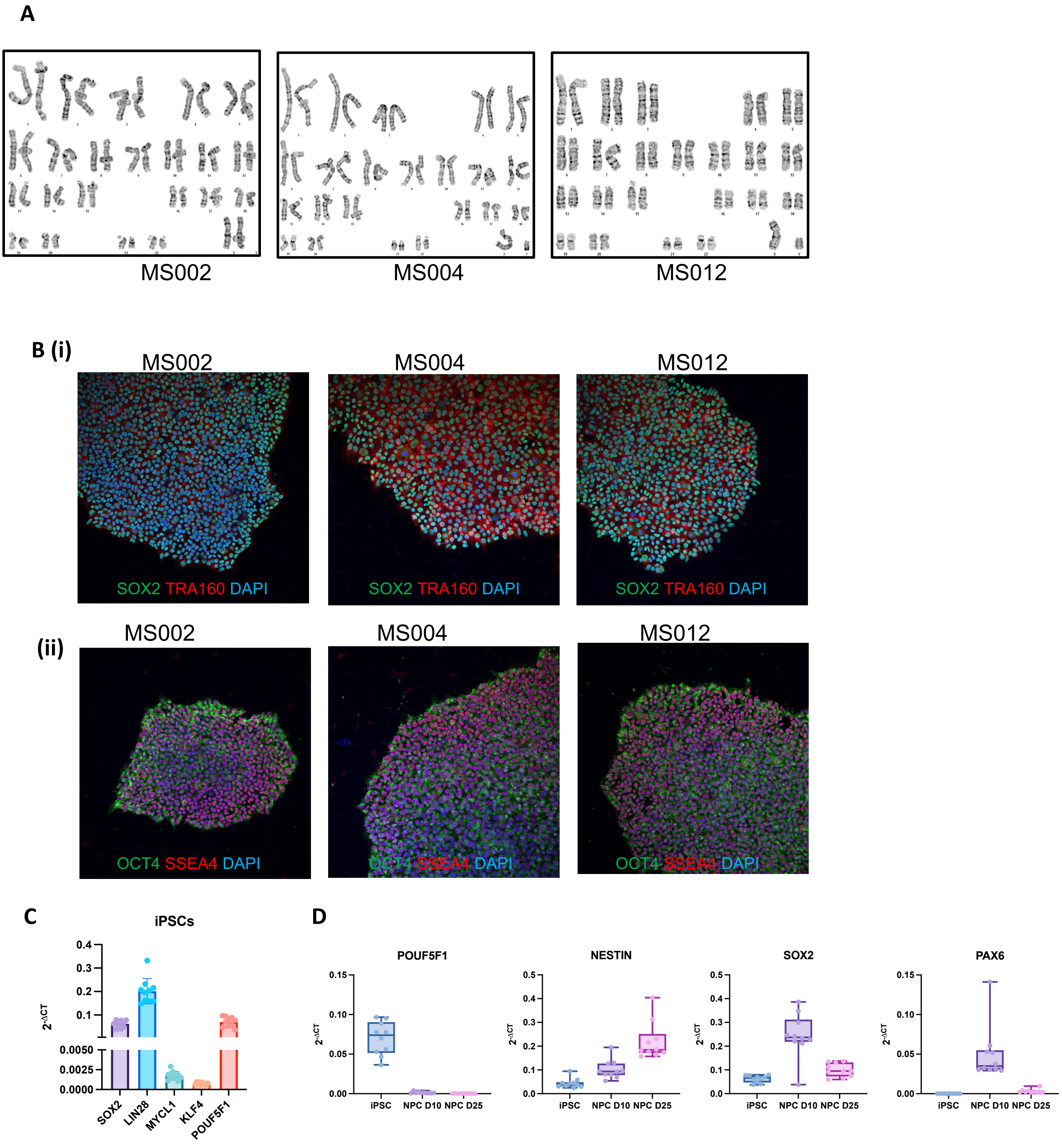


**Supplementary Figure 2. Overview of BD-FMD iPSC and NPC characterization.** (A) Karyotyping results of 3 BD-FMD patient iPSCs. (B) Representative immunofluorescence images of iPSC characterization markers SOX2, TRA160, OCT4, and SSEA4. (i) Immunofluorescence staining for SOX2 (green), TRA160 (red), and DAPI (blue). (ii) Immunofluorescence staining for OCT4 (green), SSEA4 (red), and DAPI (blue). Images taken at 10X Magnification. (C) Bar graph representing relative mRNA expression of SOX2, LIN28, MYCL1, KLF4, and POUF5F1 in iPSCs. (D) Bar graphs representing relative mRNA expression of POUF5F1, Nestin, SOX2, and PAX6 in iPSCs and NPC day 10, NPC day 25.

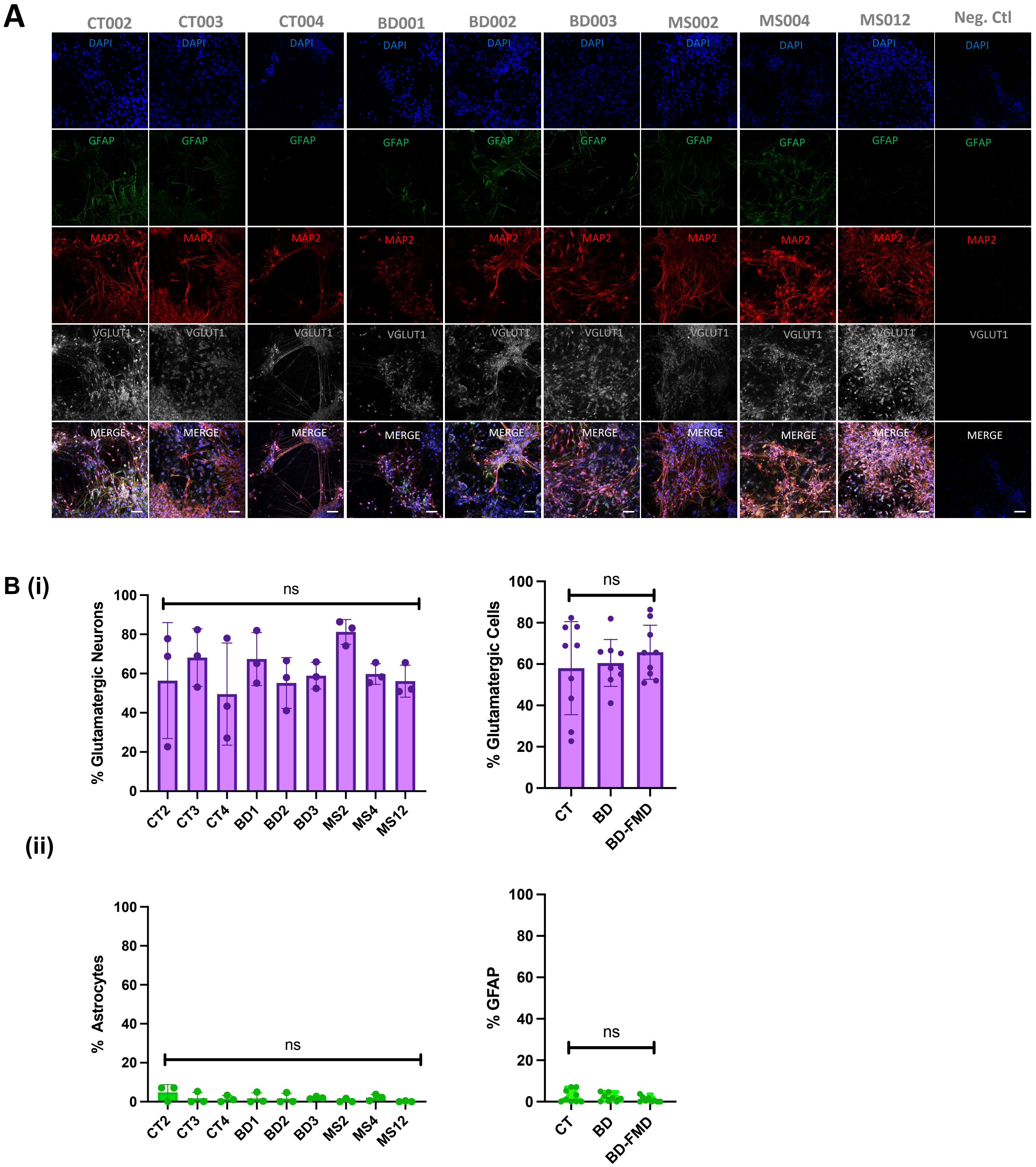


**Supplementary Figure 3.** Immunofluorescent characterization of neuronal lines at Day 50. **(A)** Representative immunofluorescent images of MAP2 (red), DAPI (blue), GFAP (green), and VGLUT1 (grey). Scale bars = 50 μm. **(B) (i)** Quantification of percent cells expressing VGLUT1 as a percentage from total DAPI in each cell line, and as a grouped comparison (CT vs BD vs BD-FMD). **(ii)** Quantification of percent cells expressing GFAP as a percentage from total DAPI in each cell line, and as a grouped comparison (CT vs BD vs BD-FMD).

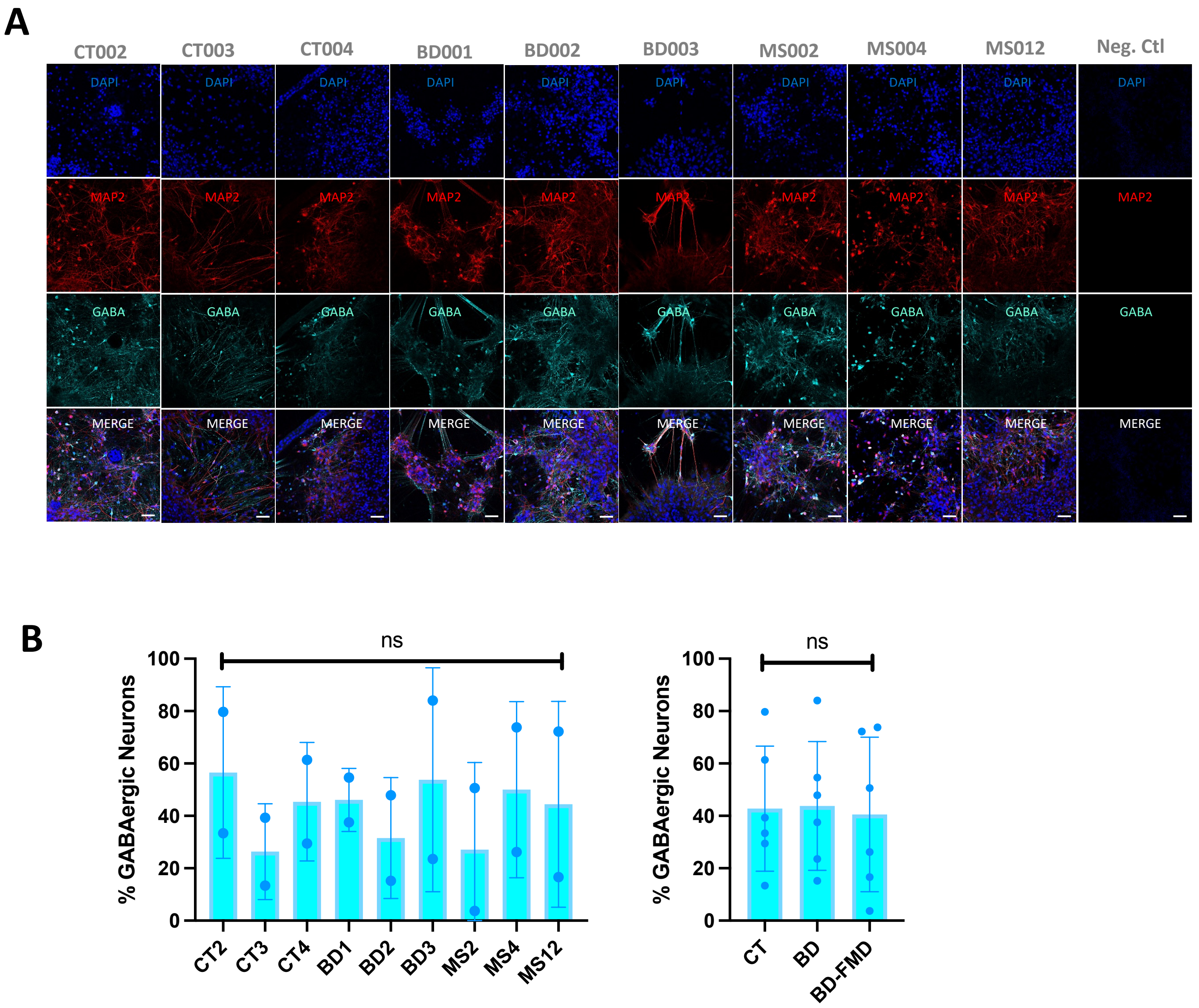


**Supplementary Figure 4. (A)** Representative immunofluorescent images of DAPI (blue), MAP2 (red), GABA (cyan). Scale bars = 50 µm. **(B)** Quantification of percent cells expressing GABA as a percentage from total DAPI in each cell line, and as a grouped comparison (CT vs BD vs BD-FMD).

**A**

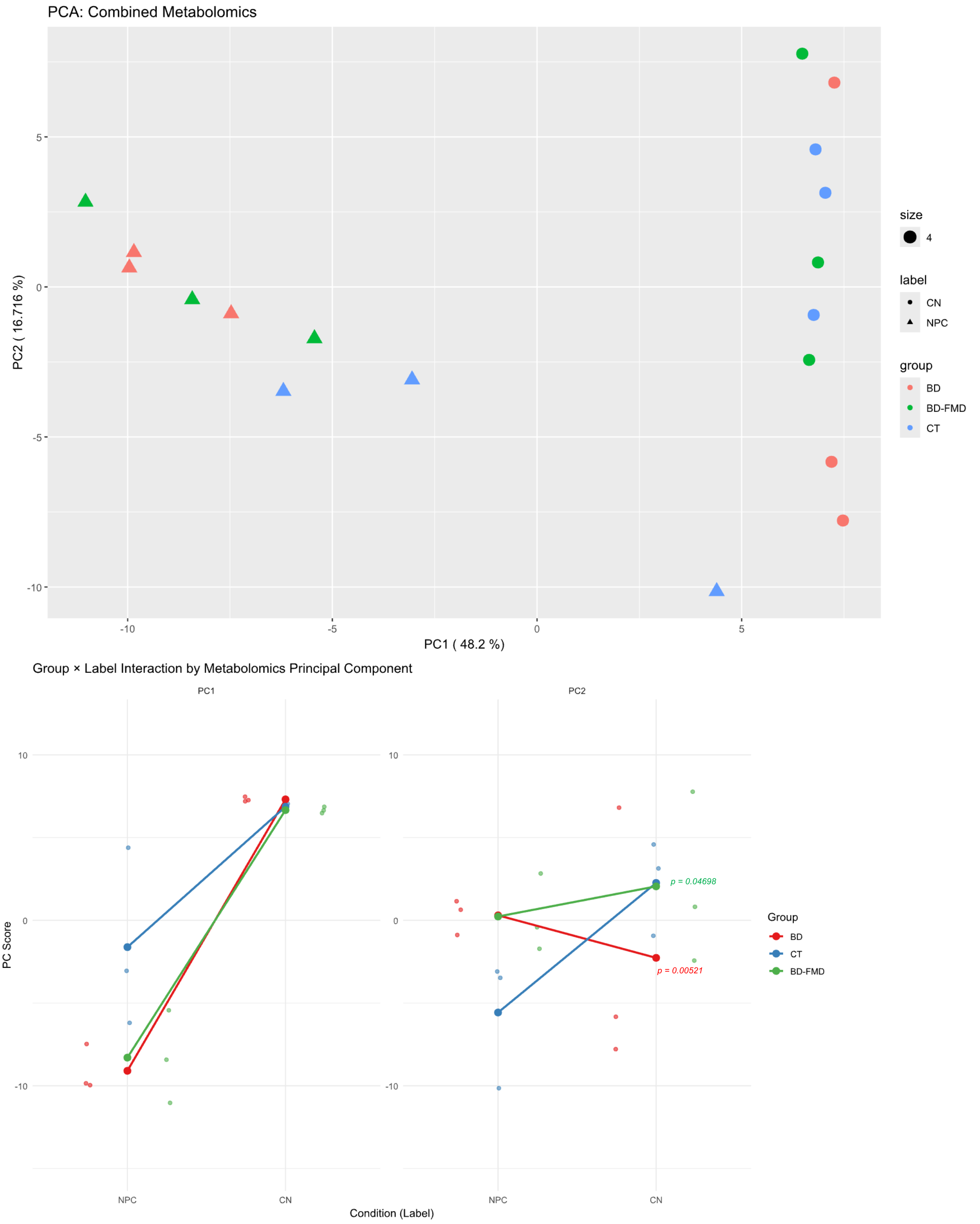


**Supplementary Figure 5**. Principle Component Analysis (PCA) of combined metabolomics data from NPCs and CNs. **(A)** PC1 (48.2%) separated samples primarily by cell type, whereas PC2 (18.1%) reflected a group × cell-type interaction. **(B)** Significant PC2 differences were observed in BD ($p = 0.00521$) and BD-FMD ($p = 0.04698$) CNs relative to controls, indicating diagnosis-specific metabolic shifts that emerge during neuronal differentiation.

**A (i)** **(ii)**

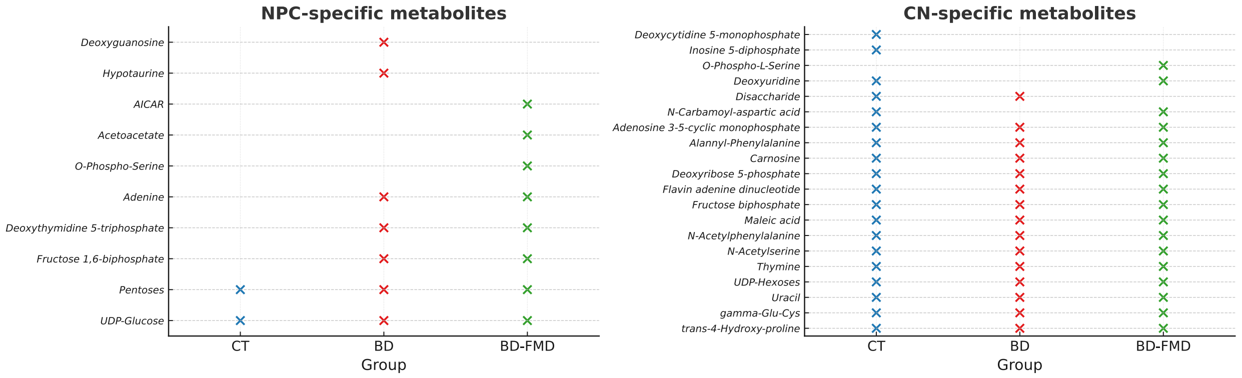


**Supplementary Figure 6. Overview of NPC and CN-specific metabolites across groups.** (A) (i) NPC-specific metabolites and (ii) CN-specific metabolites identified in each group: CT (blue), BD (red), and BD-FMD (green); n = 3 donor lines per group. A metabolite is listed as stage-specific for a group if it was detected at that stage and absent at the other stage within the same group (see Methods for detection criteria). Colored markers indicate presence in the indicated group; metabolites appearing in multiple columns are shared between groups at that stage, whereas single-column entries are group-unique. These lists provide the drivers for pathway enrichments in Figure 4 (e.g., deoxyguanosine/hypotaurine at NPC in BD; AICAR, acetoacetate, O-phospho-serine at NPC and O-phospho-L-serine at CN in BD-FMD; deoxycytidine-5′-monophosphate/inosine-5′-diphosphate at CN in CT). Abbreviations; AICAR, 5-aminoimidazole-4-carboxamide ribonucleotide.

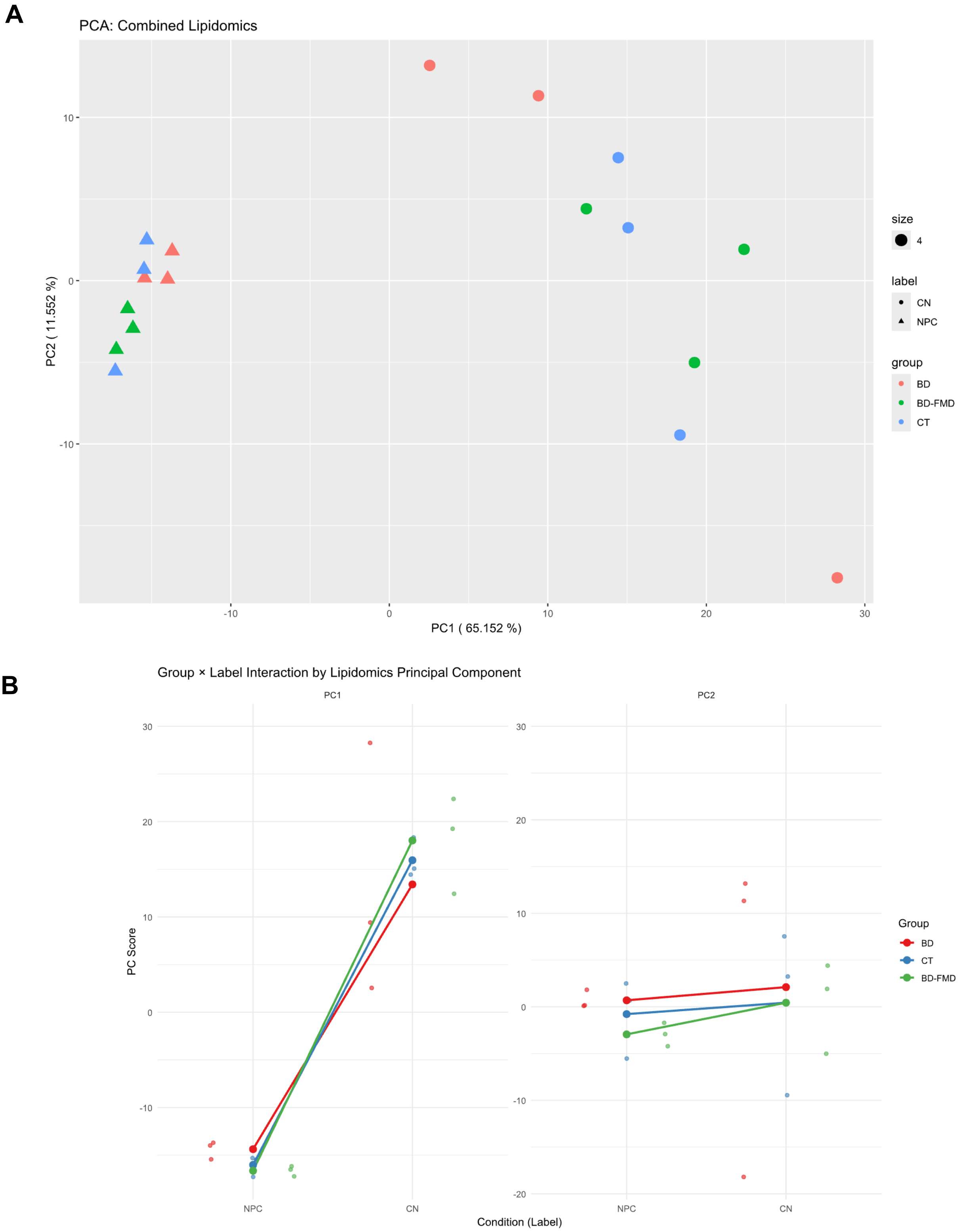


**Supplementary Figure 7**. Principle Component Analysis (PCA) of combined lipidomic data from NPCs and CNs. **(A)** PC1 (65.15%) captured cell-type–specific lipidomic variance, while PC2 (11.54%) showed no significant separation by diagnostic group. **(B)** No group and cell-type interaction reached statistical significance, suggesting that lipidomic remodeling during neuronal differentiation is not detectably altered in BD or BD-FMD lines.

### Supplementary Tables

Supplementary Table 1. Clinical Demographics and Characteristics

| Subject ID | Age | Sex | Age at Diagnosis | BD | MD | Medical history | Medication History | FMD |
|---|---|---|---|---|---|---|---|---|
| MS002 | 64 | F | 57 | YES | NO | Diabetes (type II), fibromyalgia, osteopenia, osteoarthritis | Received cognitive behavioral therapy; Lithium carbonate (750 mg/day), bupropion (300 mg/day), metformin (850mg/day) | Yes, COX Deficiency |
| MS004 | 38 | M | 26 | YES | NO | N/A | Sodium valproate (1250 mg daily) and lamotrigine (400 mg daily) | Yes, COX Deficiency |
| MS012 | 61 | M | 34 | YES | NO | N/A | N/A | Yes, COX Deficiency |
| BD001 | 38 | F | 30 | YES | NO | | 200 mg Lithium; Sertraline 100mg; Psychotherapy | NO |
| BD002 | 59 | F | 34 | YES | NO | Asthma, Osteoarthritis | Venlaflaxine 300mg; Clonazepam 2mg; Pravachol 10mg; Nexium 20mg; Vitamin D; Coloxil with Senna; Symbicort inhaler 4 pumps/day; Quetiapine 300mg | NO |
| BD003 | 36 | F | 24 | YES | NO | Complex regional pain syndrome (CRPS) | Sertraline 100m; Aripiprazole 15mg; Norflex 200mg; Dilaudid 16mg; Targin (30+15); Jurnista 8mg; Nexium 40mg | NO |
| CT002 | 42 | F | N/A | | NO | | | NO |
| CT003 | 37 | F | N/A | NO | NO | | | NO |
| CT004 | 55 | F | N/A | | NO | Hypothyroidism | | NO |

Supplementary Table 1. Clinical Demographics and Characteristics

Supplementary Table 2. Clinical Demographics and Characteristics of patients with mitochondrial disease

| Subject ID | Age of Diagnosis | Age of Death | Sex | BD | MD | Medical history |
|---|---|---|---|---|---|---|
| WG0965 | 3 MO | 5 MO | F | No | Cytochrome C oxidase Deficiency | Infantile; poor muscle development; hepatomegaly; neurological deterioration; high lactate, pyruvate, and alanine; no response to thiamine |
| WG1153 | N/A | 3.5 MO | F | No | Cytochrome C oxidase Deficiency | infantile |
| WG1210 | N/A | 7 MO | F | No | Cytochrome C oxidase Deficiency | Failure to thrive, hypotonia, severe liver failure, infantile |
| WG1669 | N/A | 3 YR | M | No | Cytochrome C oxidase Deficiency | infantile |

Supplementary Table 2. Clinical Demographics and Characteristics of patients clinically diagnosed with a Mitochondrial Disease

Supplementary Table 3. Summary of Karyotyping Results of BD-FMD and iPSC lines.

| Participant ID | Karyotyping Result |
| --- | --- |
| MS002 | 46, XX |
| MS004 | 46, XY |
| MS012 | 46, XY |

Supplementary Table 3. Summary of Karyotyping Results of BD-FMD iPSC lines. Karyotype analysis exhibited normal male (46, XY) and female (46, XX) .

Supplementary Table 4. Epi-Pluri-Score of iPSC lines

| Sample ID | DNA Methlyation percentage at 3 Cpgs | | | Epi-Pluri-Score |
|---|---|---|---|---|
| | β-value [ANKRD46] | β-value [C14orf115] | β-value [POU5F1] | |
| MS002 | 24.5 | 7.6 | 65.3 | 16.9 |
| MS004 | 23.8 | 9.6 | 82.5 | 14.2 |
| MS012 | 22.9 | 8.4 | 71.6 | 14.5 |

Supplementary Table 4. Epi-Pluri-Score DNA methylation of BD-FMD iPSC lines where Epi-Pluri-Score is calculated based on β-values represent DNA methylation at 3 Cpgs.
